# Enriching neural stem cell and pro-healing glial phenotypes with electrical stimulation after traumatic brain injury in male rats

**DOI:** 10.1101/2020.11.07.372979

**Authors:** Eunyoung Park, Johnathan G. Lyon, Melissa Alvarado-Velez, Martha I. Betancur, Nassir Mokarram, Jennifer H. Shin, Ravi V. Bellamkonda

## Abstract

Traumatic Brain Injury (TBI) by an external physical impact results in compromised brain function via undesired neuronal death. Following the injury, resident and peripheral immune cells, astrocytes, and neural stem cells (NSCs) cooperatively contribute to the recovery of the neuronal function after TBI. However, excessive pro-inflammatory responses of immune cells, and the disappearance of endogenous NSCs at the injury site during the acute phase of TBI, can exacerbate TBI progression leading to incomplete healing. Therefore, positive outcomes may depend on early interventions to control the injury-associated cellular milieu in the early phase of injury. Here, we explore electrical stimulation (ES) of the injury site in a rodent model (male Sprague-Dawley rats) to investigate its overall effect on the constituent brain cell phenotype and composition during the acute phase of TBI. Our data showed that a brief ES for 1h on day 2 of TBI promoted pro-healing phenotypes of microglia as assessed by CD206 expression and increased the population of NSCs and Nestin^+^ astrocytes at 7 days post-TBI. Also, ES effectively increased the number of viable neurons when compared to the unstimulated control group. Given the salience of microglia and neural stem cells for healing after TBI, our results strongly support the potential benefit of the therapeutic use of ES during the acute phase of TBI to regulate neuroinflammation and to enhance neuroregeneration.

**Significance Statement:** Traumatic brain injury (TBI) occurs when a head injury leads to a disruption of normal function in the brain and is a major cause of death and disability, worldwide. The authors used electrical stimulation during the acute phase of TBI, which promoted pro-healing phenotypes of microglia and increased the number of neural stem cells and Nestin^+^ astrocytes, thereby enhancing neuronal viability. These findings support further study of electrical stimulation to regulate neuroinflammation and to enhance neuroregeneration after TBI.

**Graphical Abstract:** FIGURE 1.

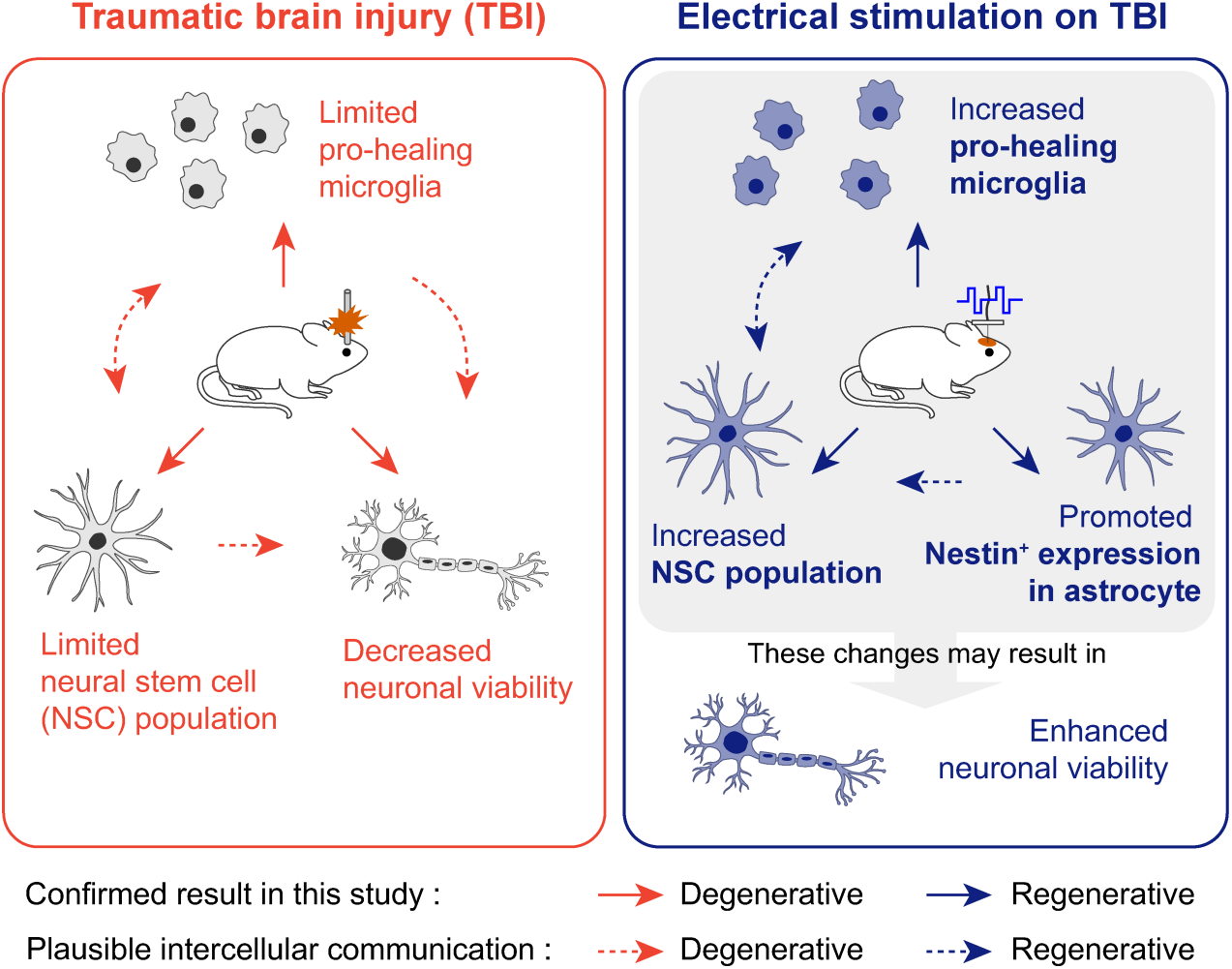

## INTRODUCTION

Traumatic Brain Injury (TBI) occurs when an external physical impact results in compromised brain function. The pathophysiological processes in TBI include physical damage to the skull and blood-brain barrier (BBB), immune cell activation, and neuronal impairment. In mild TBI– where the injury-associated disorientation or unconsciousness is shorter than 30 minutes (National Center for Injury Prevention and Control (U.S.), 2003)–the brain tissue undergoes a natural healing process with controlled hemorrhage, followed by recovery of damaged neuronal function. During the healing process, a well-orchestrated response of immune cells within the injury microenvironment is hypothesized to facilitate the recovery of a dysfunctional brain. On the other hand, severe TBI–where the injury-associated neurological states exceed 24 hours as a result of injury of substantial size or damage to a critical brain region–overwhelms one’s self-healing capability, leading to lasting disability and neural dysfunction.

At the cellular level, multiple cell types contribute to the progression and recovery of the injured brain after TBI (Simon et al., 2017). Primarily, necrotic death and structural disruptions in both neuronal and non-neuronal cells occur at the moment of impact. During the initial immune response to this damage, microglia, the resident immune cells in the brain, become activated by distressed cells in the injury microenvironment. Peripheral cells, including leukocytes (e.g., neutrophils and monocytes), T-cells, and dendritic cells, home to the injury– a process typically enhanced when an injury to the BBB is incurred. Astrocytes, one of the most abundant cells in the brain, also become activated in pathological responses. Interestingly, neural stem cells (NSCs), typically found during development as parental cells to many brain cells, are also found after brain injury. These NSCs possibly serve to replace the damaged cells and/or to alter the deleterious microenvironment toward a more neuroprotective one through secretion of favorable soluble factors (Bond, Ming, & Song, 2015). Nevertheless, in severe cases, long-lasting inflammatory reactions would lead to substantial neurodegeneration.

This chain of cellular responses is dynamic, especially in the early phase of TBI (Figure S1). For example, the early inflammatory responses, that span over several days, are mostly driven by resident microglia and infiltrated monocytes and macrophages. These cells are classified by the M1/M2 paradigm based on signatures of cytokine and chemokine secretion, reactive oxygen species production, phagocytic activity, and antigens expressions (Loane & Kumar, 2016). A few recent studies reported the temporal phenotypic switching of the immune cells from mixed M1/M2 phenotypes to M1 phenotypes in an adult rat TBI (Kumar, Alvarez-Croda, Stoica, Faden, & Loane, 2016; Simon et al., 2017; Wang et al., 2013). On the other hand, endogenous NSCs are known to be recruited to the site of cortical injury within one day after TBI while maintaining proliferation and migration from their niche for several days (Itoh, Satou, Hashimoto, & Ito, 2005; Yi et al., 2013). However, they often die before differentiating into mature neurons, possibly due to a shortage of factors necessary for their survival and differentiation (Itoh et al., 2005; Yi et al., 2013). Therefore, these acute cellular changes inevitably lead to exacerbated TBI progression and incomplete healing.

Therefore, modulation of the injury-associated cellular and inflammatory milieu during the early phase of injury has become a subject of interest. In the recovery of TBI, peripheral nerve injury, or ischemia, several anti-inflammatory and neuroprotective compounds have been shown to enhance neuronal survival and structural remodeling by either recruiting M2 immune cells or shifting their states toward M2 (X. Liu et al., 2016; Mokarram & Bellamkonda, 2011; Mokarram et al., 2017). Likewise, in case of endogenous NSCs, pharmacological growth factors or cytokines have been shown to modulate proliferation, apoptosis, migration, or differentiation lineage (Addington, Roussas, Dutta, & Stabenfeldt, 2015). In the clinical application of these pharmacological factors, however, safety issues still remain in the determination of the most effective dosage and treatment strategy that warrants no detrimental side effects.

As a therapeutic concomitant or alternative to pharmacological approaches, electrical stimulation (ES) has gained a great deal of attention over the last few decades. In neurological diseases including chronic pain, depression, Parkinson’s disease, and TBI, (Hofer & Schwab, 2019; Limousin et al., 1998; Schiff et al., 2007) the therapeutic application of ES has mainly focused on the functional recovery of neurons, to some degree neglecting glial cells in the brain (Otto & Schmidt, 2020). However, ES has the potential to modulate the physiology of many cell types in the brain (Chen, Bai, Ding, & Lee, 2019). For instance, NSCs have been shown to exhibit enhanced proliferation, differentiation, and directed migration in response to ES (Huang, Li, Chen, Zhou, & Tan, 2015; Zhu et al., 2019). Accordingly, several studies have demonstrated the applicability of ES in NSCs to elicit neurogenesis in peripheral nerve regeneration (Iwasa et al., 2019), stroke (Xiang et al., 2014), and memory dysfunction (A. Liu, Jain, Vyas, & Lim, 2015) in animal models.

In addition, recent studies have reported the efficacy of ES in the regulation of neuroinflammation. For example, in painful neuropathy by sciatic nerve transection, ES treatment ameliorated hyperalgesia by suppressing the activation of both microglia and astrocytes (Lopez-Alvarez, Cobianchi, & Navarro, 2019). In multiple sclerosis, an autoimmune disease, ES was shown to polarize macrophages toward M2 phenotypes, ultimately contributing to remyelination (McLean & Verge, 2016). Moreover, ES treatment following lipopolysaccharide exposure or spinal contusion alleviated microglial activation while improving neural activity (Hahm, Yoon, & Kim, 2015; Huffman et al., 2019).

Despite growing interests in the field, the effects of ES in TBI, on both acute neuroinflammation and surrounding tissue have not been fully elucidated. This study aims to investigate the impact of brief ES on the second day of TBI, to induce a pro-healing biochemical cascade to positively impact brain recovery by enhancing the neuroprotective responses of immune cells and endogenous NSCs, thereby enhancing neuronal viability.

## METHODS

### Surgical Procedures: TBI Induction

All animal studies were approved by the Institutional Animal Care and Use Committee (IACUC) at Duke University, and protocols were performed following the Guide for the Care and Use of Laboratory Animals published by the National Institute of Health (NIH). A total of 24 eight-week-old male SAS Sprague-Dawley rats weighing 250-300 g were obtained from Charles Rivers Labs (Crl:CD(SD), Strain code: 400, RRID:RGD_734476). All animals received a craniotomy and controlled cortical impact (CCI), following procedures used in prior work by our group (Betancur et al., 2017). Briefly, a longitudinal incision was made, and a 5 mm craniotomy was performed 0.5 mm anterior to bregma and 0.5 mm lateral from the sagittal suture. After removing the bone flap, a 3 mm diameter injury with a depth of 2 mm with a speed of 4 m/s was made, located at the center of the craniotomy. The severity of CCI is considered to be moderate to severe. The injury site was covered entirely with a BloodStop Hemostatic Gauze (Life Science Plus). The skin flaps were subsequently sutured together to close the wound, and triple antibiotic cream was layered on top of the sutured skin. The animals received a Buprenorphine (1 mg/kg) injection and were allowed to recover in a new, clean cage. In all surgical procedures, each rat was anesthetized using 2 % isoflurane gas with 100 % oxygen level, and placed on a heated pad to maintain its body temperature at 37 °C. The head was held in a stereotaxic frame (David Kopf Instruments, CA) with the snout placed into a nose cone to deliver the aforementioned level of surgical anesthesia.

### Surgical Procedures: Electrode Implantations and Application of the ES

Two days post-TBI, animals were randomly assigned to the sham and ES groups (8, 7, and 9 animals were chosen as the TBI control, sham, and ES group, respectively) (Figure 1a,b). For the sham group, an electrode was implanted without ES stimulation, whereas rats in the ES group received electrical stimulation through the implanted electrode.

**FIGURE 1.**
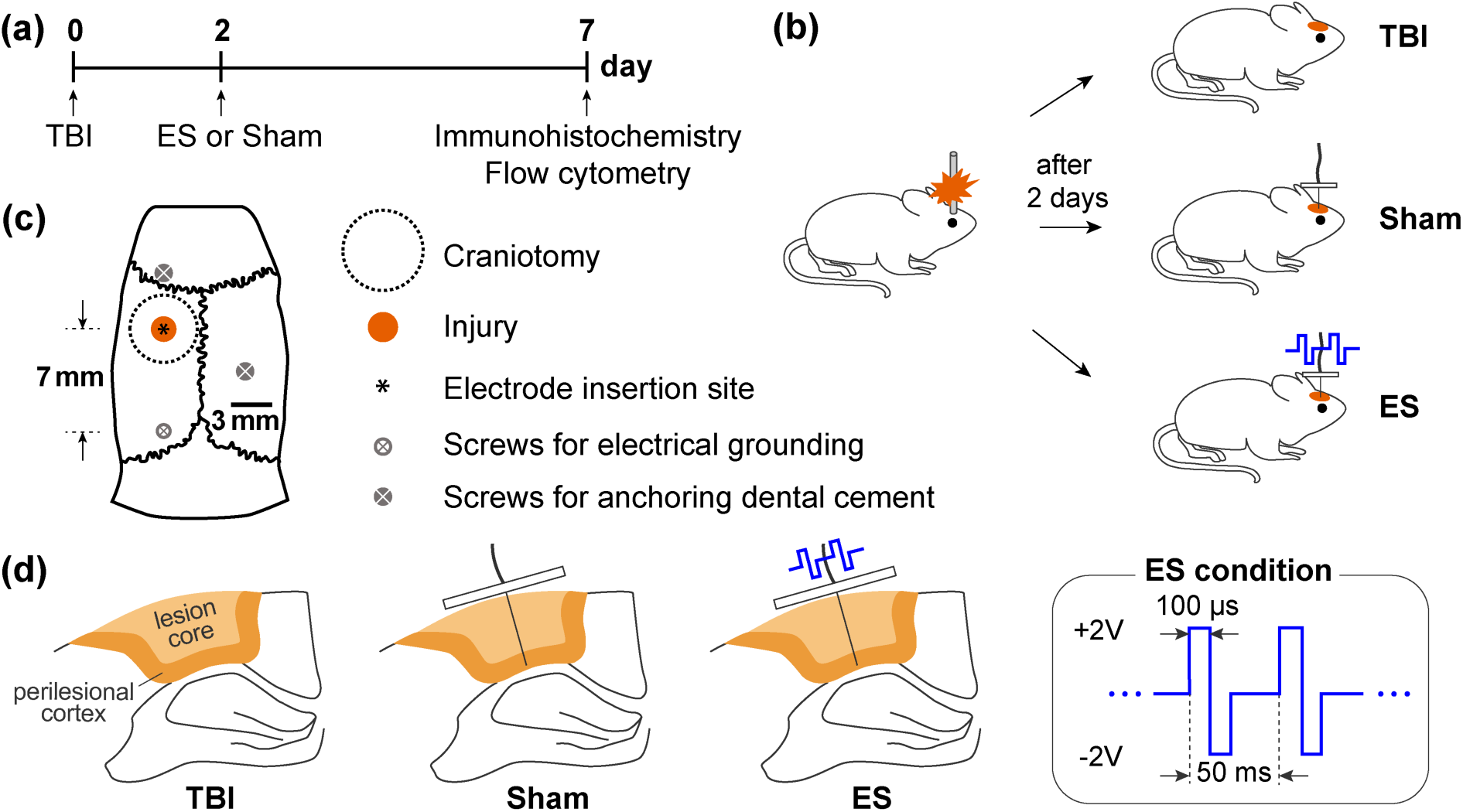
Experimental design. (a) Timing of experiments, including traumatic brain injury (TBI) induction, sham and electrical stimulation (ES) operations, and analysis. (b) All animals received a controlled cortical impact (CCI) for the TBI. In two days post-TBI, rats were randomly divided into three groups: untreated TBI (untreated control group), sham (TBI with an implanted electrode without ES), and ES (TBI with applied electrical stimulation through the implanted electrode) groups. (c) Schematic of contusion region induced by CCI system and the position of the inserted electrode on a rat skull. Screws used to anchor the dental cement or connect the electrical grounding. (d) Schematic of the coronal section of perilesional brain to show an inserted electrode and ES conditions.

We designed an implantable ES device, made of a 200 µm diameter platinum microwire electrode (Omega Engineering, Cat# SPPL-008), 255 µm Silicon/Copper hookup wire, and a polydimethylsiloxane (PDMS; Ellsworth Adhesives, Sylgard 184) block (Figure S2). The assembled ES devices were sterilized by immersion in 70 % ethanol within a UV light sterilizer chamber for one hour. A Germinator dry bead sterilizer was also used prior to the implantation. The animals for the sham and ES groups were prepared by placing them under surgical anesthesia as described above. The incision area was sanitized using ethanol and chlorhexidine, and the sutures were removed to release the skin flaps. An electrode was implanted in the injury epicenter to a depth of 2 mm in order to stimulate only the injured cerebral cortical region (Figure 1c,d). Three stainless steel screws were implanted in the skull to assist in affixing the device in place: the first at the position 1.5 mm anterior to the coronal suture, the second 2 mm lateral from the sagittal suture on the contralateral side, and the third (also used for connecting to electrical ground) 6 mm posterior to the electrode. The ES devices, including all screws, were covered with UV curing dental cement. For sham animals, the skin flaps were sutured, and the animal was allowed to recover as described above. In the ES group, after CCI and electrode implantation, the electrode and ground leads were connected to a function generator (Rigol, DG1022). ES was delivered as a rectangular, symmetric, biphasic, pulses (4 V peak-to-peak (V_pp_), 20 Hz frequency, 100 microseconds pulse duration) for one hour (Figure 1d). 4 V_pp_ was chosen as it was the maximal stimulation voltage that did not result in animals twitching during the stimulation. Note, the details of these conditions should the optimized for different animals or experimental conditions/models. Here, rectangular, symmetric, biphasic, pulsed ES was chosen to minimize persistent charge gradients, and deleterious electrodic effects (Merrill, Bikson, & Jefferys, 2005). After ES treatment, rats were also allowed to recover like sham animals.

### Neural Tissue Preparation and Immunohistochemistry

Seven days post-injury, animals were sedated using 5 % isoflurane gas and transcardially perfused with 200 mL phosphate-buffered saline (PBS) (pH 7.4) followed by 100 mL 20 % sucrose in PBS. The brains were extracted and cut at the epicenter of the lesion using a rat brain matrix in the coronal plane (Ted Pella Inc., CA). Two half-sections submerged in optimal cutting temperature compound (Tissue Tek, Miles Inc., IN) were frozen in liquid nitrogen and stored at −80 °C. The frozen brains were sectioned at 12 µm thickness using a cryostat (LeicaBiosystems, IL) and then collected onto glass slides: placing three 12 μm sections from the rostral side of the injury and three from the caudal side in each slide, and collecting ten slides per animal. Tissues were fixed with 100 % ethanol for 1 minute and then washed with PBS for 5 minutes, three times. Slides were kept in −20 °C before immunohistochemical staining and taken out before the staining to reach room temperature.

Immunohistochemistry was performed following previously described methods and controls matching previously published and appropriate patterns of stained cellular morphology and distribution (Betancur et al., 2017). Slides were washed in PBS (5 minutes, three times), and permeabilized with 0.05% Triton X-100 in PBS (1 minute), repeated three times. Slides were blocked with a blocking solution (0.5 % Triton X-100 in PBS and 4 % goat serum) for one hour. Primary antibodies diluted in blocking solution were added and slides were kept overnight at 4 °C. Primary antibodies used for this study were rabbit anti-NeuN (1:400, Millipore Sigma, Cat# MABN140, RRID:AB_2571567), mouse anti-Nestin (1:500, Novus Biologicals, Cat# MAB2736, RRID:AB_2282664), rabbit anti-Glial fibrillary acidic protein (GFAP; 1:1000, Dako, Cat# Z033429, RRID:AB_10013382), mouse anti-CD68 (ED1; 1:400, BioRad, Cat# MCA341R, RRID:AB_2291300), and rabbit anti-Mannose receptor (CD206; 1:500, Abcam, Cat# ab64693, RRID_AB1523910). Secondary antibodies diluted in PBS were added to samples for one hour at room temperature: Alexa Fluor 488-conjugated goat anti-rabbit IgG (1:400, Abcam, Cat# ab150081, RRID:AB_2734747), and Alexa Fluor 594-conjugated goat anti-mouse IgG (1:220, Abcam, Cat# ab150116, RRID:AB_2650601). The prepared slides were then immersed in DAPI (1 μg/mL, Sigma Aldrich, Cat# D9542) solution for 20 minutes. Between each staining step, samples were washed thoroughly three times with permeabilization solution and PBS sequentially for 1 min and 5 mins, respectively. Slides were covered with coverslips supplemented with Fluoromount-G (Southern Biotech, AL).

### Microscopy

Axio Observer 7 (Carl Zeiss, Germany) equipped with an Axiocam 702 mono camera or equipped with an Axiocam 305 color camera was used for imaging immunohistochemical tissues or Nissl-stained tissues, respectively. Confocal z-stack images were taken with LSM 880 (Carl Zeiss, Germany).

### Image Quantification

Quantitative analysis of images was done by tools in ImageJ (FIJI, RRID:SCR_002285). For analysis, two brain sections were used, one within 1 mm rostral and one within 1 mm caudal to the lesion epicenter. In particular, before analyzing the intensity profiles, area, and degree of colocalization for specific fluorophore^+^ pixels, image intensity processing was performed. For the intensity profile analysis, images were processed with the Subtract Background tool to fix an uneven background then measured for intensity along the ROI line using the Plot Profile tool with a line width of 300 pixels. For fluorophore^+^ area and colocalization analysis, background correction for images was performed using the Subtract Background tool, and then thresholding used to make fluorophore^+^ pixels white and all other pixels black. Total fluorophore^+^ area was measured using the Analyze Particles tool. In this study, Colocalization Threshold tool was used to obtain two Mander’s split coefficients (M1 and M2: the colocalization coefficient relative to channel 1 and channel 2), and % of colocalized pixels (%Ch1 Vol and %Ch2 Vol: the percentage of the total number of colocalized pixels for each channel above their respective thresholds) (Zinchuk, Zinchuk, & Okada, 2007). The colocalized area was calculated by multiplying the channel_1_^+^ area and %Ch1 Vol (the same value as channel_2_^+^ area multiplied by the %Ch2 Vol). Because we measured the split coefficients using threshold-adjusted images, we minimized in quantification error due to background noise.

### Flow Cytometry

Flow cytometry was performed to analyze the phenotypes of immune cells in the ipsilateral brain following a modified version of a previously published protocol (Posel, Moller, Boltze, Wagner, & Weise, 2016). The brain was extracted and placed into a stainless-steel coronal rat brain matrix (Zivic Instruments, Cat# BSRAS001-1), making coronal cuts to obtain 5 mm thick slices (brain slices at 1 - 6 mm posterior to bregma). The 5 mm thick brain slices were transferred into a plastic petri dish, placing the coronal plane on the dish surface, and the brain slices that contained the injured tissue of ipsilateral-cortex and hippocampus were dissected (5 × 5 mm^2^). The dissected tissue samples were then dissociated into single-cell suspension. Cells were suspended in flow cytometry staining buffer (BD Biosciences) and treated with Fc-receptor blocker (2.5 μ g/mL, anti-mouse CD16/32; BioLegend, Cat# 101301, RRID: AB_312800) for 20 minutes at 4 °C in the dark. For the cell surface staining, fluorophore-conjugated primary antibodies, CD45-Pe/Cy7 (1 μ g/mL, BD Biosciences, Cat# 561588, RRID:AB_10893200) was used. CD206 (1 μ g/mL, Abcam, Cat# ab64693, RRID:AB_1523910) antibody was used with 488-conjugated secondary antibody (1 μg/mL, Abcam, Cat# ab150081, RRID:AB_2650601). Cells were stained by these antibodies for 30 minutes at 4 °C in the dark. Following thorough washing, flow cytometry was performed with a Novocyte 2060 flow cytometer and analyzed using FlowJo software (TreeStar, Inc., OR, RRID:SCR_008520). The specificity of the signals of antibodies against specific antigens was determined by performing a control experiment using compensation beads (Invitrogen, Cat# 01-1111-42).

### Graphing and Statistics

Treatment groups were randomly assigned following TBI induction. For sectioning, 10 slides per rat were obtained, and slides were randomly assigned for immunohistochemistry. For flow cytometry, the experiments were performed at once. One of the ES samples was found to be an outlier (as analyzed by Dixon’s Q-test; 4 of 6 stained values were found to be an outlier at >90% confidence for this replicate), thus this rat was excluded. Outlier analysis was not performed on immunohistochemistry data. A power analysis was not performed a priori.

Prior to the statistical analysis, a Shapiro-Wilk normality test was performed for all data sets, and a statistical analysis method was chosen accordingly from: Kruskal Wallis 1-way ANOVA with Dunn’s post-hoc, ANOVA 2-way with Tukey’s post-hoc, or Welch ANOVA with Dunnett’s post-hoc. All graphs and statistical analyses were done with Prism 8 (Graphpad Inc., RRID:SCR_002798). The statistical methods used are reported for each result, in place. An *α* of 0.05 was used to determine significance. All graphs are depicted using Tukey method box and whiskers unless otherwise specified. All data are reported as mean ± standard deviation unless otherwise specified.

## RESULTS

### ES treatment increased the number of CD206^+^ cells in the perilesional cortex

Neuroinflammation in TBI-associated pathological processes is known to have a substantial influence on TBI outcomes (McKee & Lukens, 2016; Simon et al., 2017). Especially during the acute phase of TBI, a large number of immune cells, including the resident microglia and infiltrated monocytes and macrophages, play critical roles in inflammatory responses in brain, where their functional phenotypes reflect disease progression (Simon et al., 2017).

To identify the functional states of monocytes, macrophages, and microglia following TBI, immunohistochemistry was performed to measure the expressions of CD68, a marker for monocyte/macrophage/microglia, and CD206, a marker for M2 phenotype (Figure 2). Three experimental groups were tested, namely TBI (untreated control group), sham (TBI with an implanted electrode without ES), and ES (TBI with applied electrical stimulation through the implanted electrode) groups (Figure 1b). In a qualitative comparison, the distribution of CD68^+^ cells were similar in all experimental groups. CD68^+^ cells were found in the perilesional cortex, including the core of the cortical lesion, and along the corpus callosum, but rarely in the hippocampus (Figure 2b). On the other hand, CD206^+^ cells, representing M2 phenotype cells, were found only from the cortex to the hippocampus, in lower quantities than CD68^+^ cells (Figure 2c). In addition, CD206^+^ cells exhibited a distinct spatial distribution at the perilesional cortex upon ES treatment. In both untreated TBI and sham groups, CD206^+^ cells only appeared near the cortical surface of the injury whereas those in the ES group were observed in areas deeper into the cortex.

**FIGURE 2.**
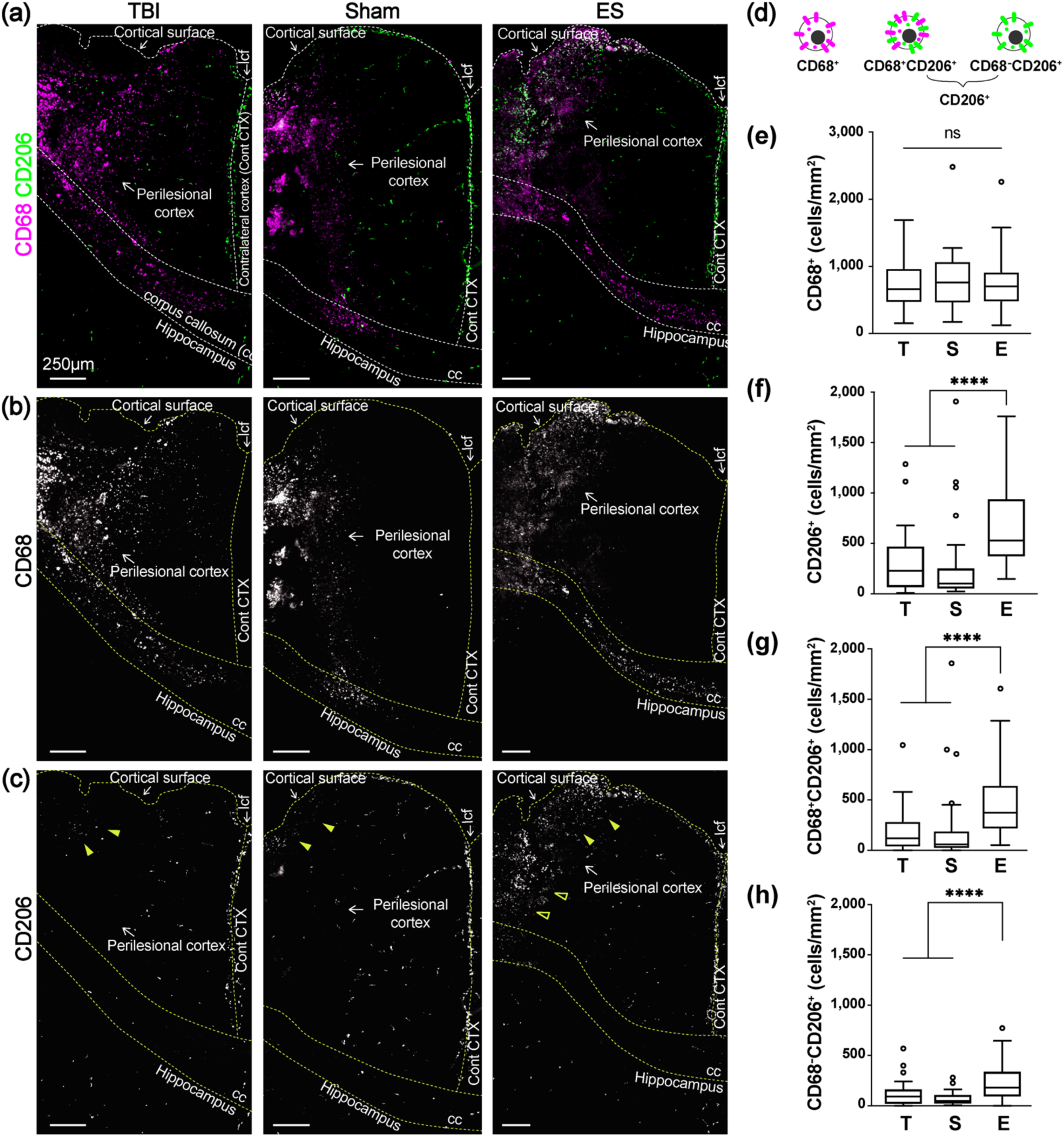
Effect of ES treatment on CD68 and CD206 expression in perilesional cortex at 7 days post-TBI. (a) Representative low magnification images of CD68 (magenta) and CD206 (green) staining from the ipsilateral cortex to the hippocampus. (b) CD68^+^ cells mostly appeared at the perilesional cortex and corpus callosum (cc), but not in the hippocampus. CD68^+^ cells were shown to be distributed similarly in all experimental groups. (c) CD206^+^ cells appeared relatively less than CD68, and they were observed at the longitudinal cerebral fissure (lcf), ipsilateral cortex, and hippocampus. Within the perilesional cortex, CD206^+^ cells were observed near the cortical surfaces in all experimental groups (filled arrowheads). On the other hand, CD206^+^ cells at the deeper cortex were only observed in the ES group (empty arrowheads). (d) Illustration of three subtypes of immune cells based on CD68 and CD206 expression. (e-h) The number of subtype cells across the entire perilesional cortex. T, S, and E on the x-axis represent untreated TBI, sham and ES groups. (e) The number of CD68^+^ monocyte/macrophages/microglia did not exhibit a significant difference by treatment. In the entire perilesional cortex, the number of (f) CD206^+^ cells, (g) CD68^+^CD206^+^ cells, and (h) CD68^-^CD206^+^ cells significantly increased after ES treatment. Data are represented as mean ± SD. N = 4, 3, 6 animals with two tissue slices analyzed for untreated TBI, sham, ES; **** *p* < 0.0001, Kruskal-Wallis one-way ANOVA with Dunn’s post-hoc analysis.

For quantitative analysis, we analyzed the density of three subtypes of immune cells, namely CD68^+^, CD68^+^CD206^+^, and CD68^-^CD206^+^ across the entire perilesional cortex (Figure 2d and Table 2).The data indicated no change in the number of CD68^+^ cells with ES treatment (*p* = 0.7386, *Kruskal-Wallis statistic* = 0.6059, *df* =2) (Figure 2e), whereas the number of total CD206^+^ cells (CD68^+^CD206^+^ and CD68^-^CD206^+^ cells) significantly increased by approximately 2.3 folds with ES compared to untreated TBI and sham operations (*p* < 0.0001, *Kruskal-Wallis statistic* = 42.67,*df* = 2) (Figure 2f).However, the overall number of CD206^+^ cells were fewer than that of CD68^+^ cells.The ES-induced increase in CD206 expression was confirmed among CD68^+^CD206^+^ (Figure 2g) and CD68^-^CD206^+^ (Figure 2h) groups,having more than 2 folds higher number in ES compared to untreated TBI and sham groups (*p* < 0.0001, *Kruskal-Wallis statistic* = 40.53, *df* = 2 for CD68^+^CD206^+^; *p* < 0.0001, *Kruskal-Wallis statistic* = 35.17, *df* = 2 for CD68^-^CD206^+^).

**Table 1.** Information on the antibodies used in this study.

| **Antibody** | **Immunogen Structure** | **Manufacturer** | **#cat/RRID** | **Species** | **Concentration** |
| --- | --- | --- | --- | --- | --- |
| NeuN | Recombinant protein corresponding to mouse NeuN (further details proprietary) | Millipore-Sigma | MABN140/AB_2571567 | Rabbit monoclonal IgG | 1:400 |
| Nestin | E. coli-derived recombinant rat Nestin Met544-Glu820 (Gly756Asp, Ile758Met, Arg572Lys, Ala574Pro, Ile802Met, Arg816Lys) Accession # EDM00749 | Novus Biologicals | MAB2736/AB_2282664 | Mouse monoclonal IgG2a | 1:500 |
| GFAP | GFAP isolated from cow spinal cord | Dako (Agilent) | Z033429-2/AB_10013382 | Rabbit polyclonal Ig fraction | 1:1000 |
| CD68 | Rat spleen cells | BioRad | MCA341R/AB_2291300 | Mouse monoclonal IgG1 | 1:400 |
| CD206 | Synthetic peptide conjugated to KLH derived from within residues 1400 to the C-terminus of Human Mannose Receptor | Abcam | Ab64693/AB_1523910 | Rabbit polyclonal IgG | 1:500 |
| Alexa Fluor 488 anti-rabbit | Details not available from manufacturer | Abcam | Ab150081/AB_2734747 | Goat polyclonal IgG | 1:400 |
| Alexa Fluor 594 anti-mouse | Details not available from manufacturer | Abcam | Ab150116/AB_2650601 | Goat polyclonal IgG | 1:220 |
| CD16/32<br>(Fc-Receptor) | Sorted pre-B<br>cells | Biolegend | 101301/AB_312800 | Rat IgG2a,<br>$\lambda$ | 2.5 $\mu\text{g/mL}$ |
| PE-Cy7<br>anti-CD45 | Leukocyte<br>Common<br>Antigen-<br>enriched<br>Glycoprotein<br>Fraction from<br>Wistar Rat<br>Thymocytes | BD<br>Biosciences | 561588/AB_10893200 | Mouse<br>BALB/c<br>IgG1K | 1 $\mu\text{g/mL}$ |

**Table 2.** The number of CD68^+^, CD206^+^, CD68^+^CD206^+^, and CD68^-^CD206^+^ cells in the perilesional cortex. Data are reported as Mean ± SD. N = 4, 3, 6 animals with two tissue slices analyzed for untreated TBI, sham, ES.

| Phenotype | ROI | Number of cells (cells/mm<sup>2</sup>) |  |  | *p*-values |  |  |
| --- | --- | --- | --- | --- | --- | --- | --- |
|  |  | TBI | sham | ES | TBI vs. sham | TBI vs. ES | sham vs. ES |
| CD68<sup>+</sup><br>(Figure 2e) | entire perilesional cortex | 711.1 ± 351.1 | 801.1 ± 458.5 | 740.9 ± 342.6 | *p* > 0.9999 | *p* > 0.9999 | *p* > 0.9999 |
| CD206<sup>+</sup><br>(Figure 2f) |  | 295.0 ± 276.6 | 297.6 ± 438.4 | 683.3 ± 396.3 | *p* > 0.9999 | ***p* < 0.0001** | ***p* < 0.0001** |
| CD68<sup>+</sup>CD206<sup>+</sup><br>(Figure 2g) |  | 182.3 ± 207.0 | 227.9 ± 420.9 | 456.0 ± 304.0 | *p* > 0.9999 | ***p* < 0.0001** | ***p* < 0.0001** |
| CD68<sup>-</sup>CD206<sup>+</sup><br>(Figure 2h) |  | 112.6 ± 113.5 | 69.7 ± 66.8 | 227.3 ± 165.4 | *p* = 0.3664 | ***p* < 0.0001** | ***p* < 0.0001** |

Kruskal-Wallis one-way ANOVA with Dunn's post-hoc analysis.

### Distribution of CD206^+^ cells shifted deeper into the perilesional cortex after ES treatment

To analyze the differential spatial distribution of three subtypes of immune cells, the injured cortex was divided into two regions, ROI_1_ and ROI_2_, the regions of 0-1 mm and 1-2 mm from the cortical surface, respectively (Figure 3a,b, and Table 3).The number of CD68^+^ cells in the two ROIs were not significantly different in all groups (two-way ANOVA; interaction, *F*_*(2, 129)*_ = 2.263, *p* = 0.1082; region, *F*_*(1, 129)*_ = 0.3986, *p* = 0.5289; treatment, *F*_*(2, 129)*_ = 0.5577, *p* = 0.5739) (Figure 3c), whereas CD206^+^ subtype cells were both region- and treatment-dependent (two-way ANOVA; for CD206^+^ cells, interaction, *F*_*(2, 129)*_ = 0.9397, *p* = 0.3934; region, *F*_*(1, 129)*_ = 22.09, *p* < 0.0001; treatment, *F*_*(2, 129)*_ = 20.81, *p* < 0.0001; for CD68^+^CD206^+^ cells, interaction, *F*_*(2, 129)*_ = 1.946, *p* = 0.1471; region, *F*_*(1, 129)*_ = 14.39, *p* = 0.0002; treatment, *F*_*(2, 129)*_ = 13.04, *p* < 0.0001; for CD68^-^CD206^+^ cells, interaction, *F*_*(2, 129)*_ = 0.3842, *p* = 0.6818; region, *F*_*(1, 129)*_ = 17.26, *p* < 0.0001; treatment, *F*_*(2, 129)*_ = 19.02, *p* < 0.0001) (Figure 3d-f). In untreated TBI and sham groups, CD206^+^ cells were predominantly observed within ROI_1_ and this level within ROI_2_ was significantly decreased by more than 2.5 folds (Figure 3d).While the number of CD206^+^ cells within ROI_1_ for the ES group was similar to that in the TBI and sham groups, interestingly, within ROI_2_, ES treatment remarkably increased the number of CD206^+^ cells, which corresponded to the amount found in ROI_1_ and was statistically significant compared to other groups. CD68^+^CD206^+^ cells exhibited similar tendency to CD206^+^ cells (Figure 3e). There were fewer CD68^-^CD206^+^ cells than CD68^+^CD206^+^ cells in both ROIs for all experimental groups, however, a significant increase in CD68^-^CD206^+^ cells due to ES was observed in both ROI_1_ and ROI_2_ (Figure 3f).

**Table 3.**
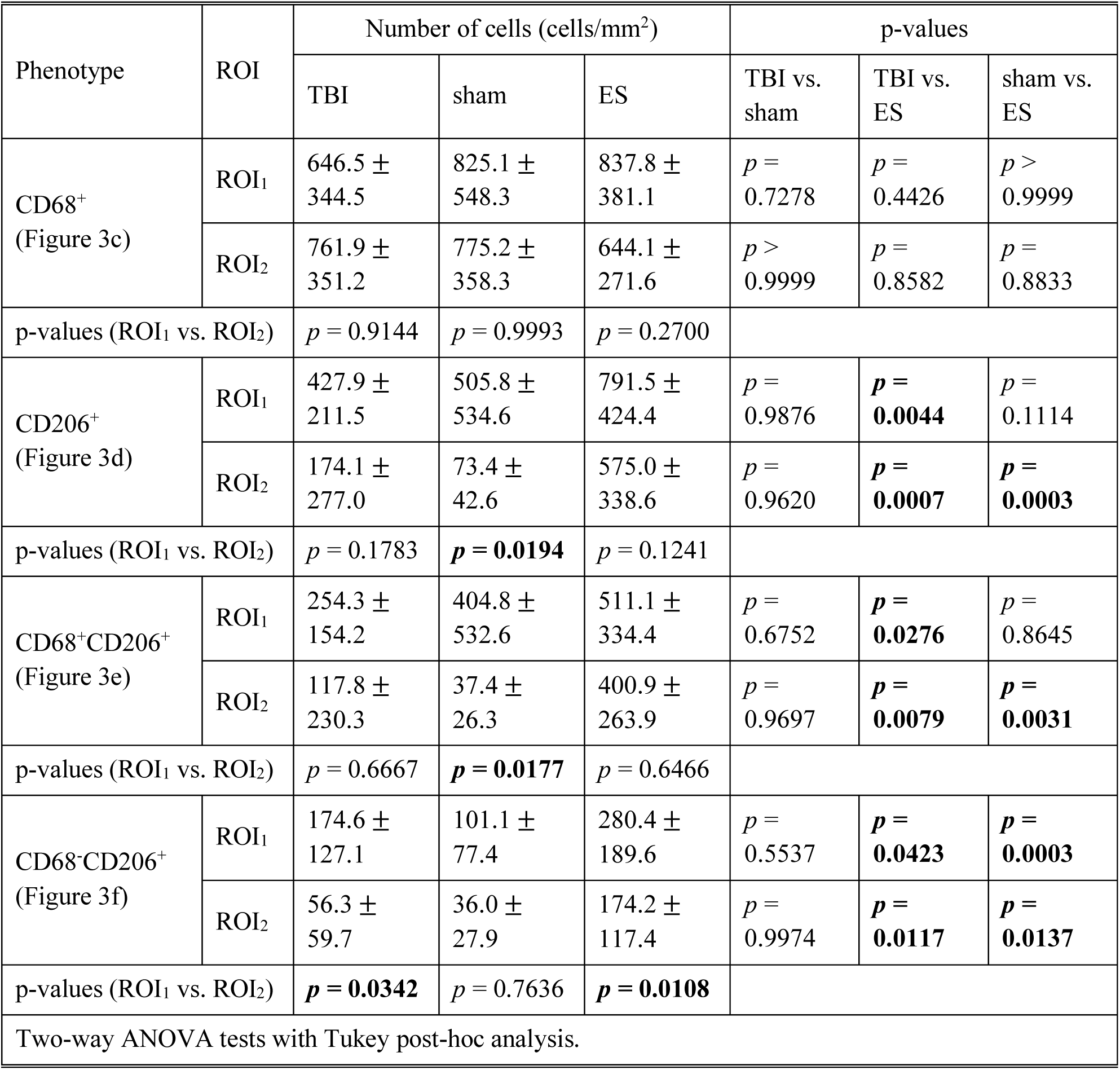
Region- or treatment-dependent expression in CD68^+^, CD206^+^, CD68^+^CD206^+^, and CD68^-^CD206^+^ cells. Data are reported as Mean ± SD. N = 4, 3, 6 animals with two tissue slices analyzed for untreated TBI, sham, ES.

**FIGURE 3.**
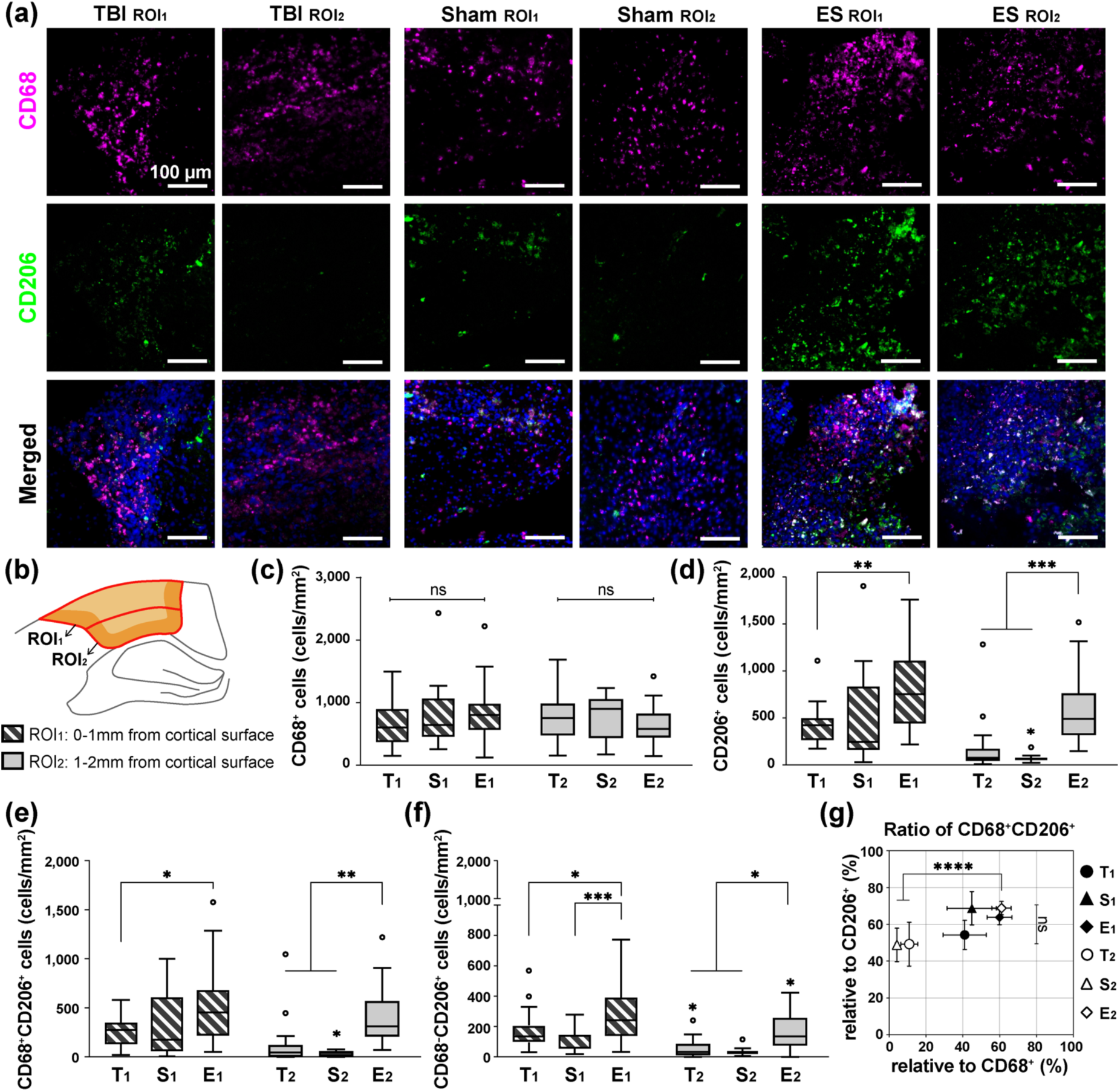
Region- or treatment-dependent expression in CD68 and CD206 at 7 days post-TBI. (a) Representative fluorescence images of CD68 (magenta), CD206 (green), and DAPI (blue) in ROI_1_ and ROI_2_. The detailed description of ROIs is included in the same Figure b. (b) Within the perilesional region, the analyzing window was divided in two: ROI_1_ and ROI_2_, positioned within 0-1 and 1-2 mm from the cortex, respectively. T, S, and E on the x-axis (c-f) or next to symbol (g) represent untreated TBI, sham and ES groups and a lower subscript 1 and 2 represent ROI_1_ and ROI_2_. (c) The number of CD68^+^ cells was not region- or treatment-dependent. (d) The number of CD206^+^ cells. Within the ROI_1_, ES treatment increased the number of CD206^+^ cells compared to untreated TBI, but not significant with sham. Instead, within the ROI_2_, CD206^+^ cells significantly increased by the ES treatment compared to TBI and sham groups. (e) The number of CD68^+^CD206^+^ cells showed a similar trend with CD206^+^ as shown in Figure d. (f) The number of CD68^-^CD206^+^ cells apparently increased by ES treatment compared to untreated TBI and sham groups in both ROI_1_ and ROI_2_. (g) The proportion of the CD68^+^CD206^+^ cells relative to the CD68^+^ (x-axis) and CD206^+^ (y-axis) cells. Treatment did not affect the proportion of CD68^+^CD206^+^ cells to CD68^+^ cells in ROI_1_, but ES treatment effectively increased those values in ROI_2_. However, there was no significant difference in the proportion of CD68^+^CD206^+^ cells to CD206^+^ cells. Data are represented as mean ± SD for Figure c-f and mean ± SEM for Figure g. N = 4, 3, 6 animals with two tissue slices analyzed for untreated TBI, sham, ES. 3-5 images were taken from each ROI; * *p* < 0.05, ** *p* < 0.01, *** *p* < 0.001, **** *p* < 0.0001, Two-way ANOVA tests with Tukey post-hoc analysis.

We then evaluated the proportion of CD68^+^CD206^+^ cells relative to CD68^+^ and CD206^+^cells in ROI_1_ and ROI_2_ (Figure 3g and Table 4).Within ROI_1_, the proportions of CD68^+^CD206^+^ cells among CD68^+^ cells of all experimental groups was not statistically different, ranging from 40 to 60 %, however, that proportion within ROI_2_ was maintained in the ES group only (61.0 ± 5.3 %) where TBI (11.0 ± 4.6 %) and sham (4.3 ± 0.8 %) groups significantly decreased compared to ROI_2_ for ES (two-way ANOVA; interaction, *F*_*(2, 45)*_ = 4.137, *p* = 0.0224; region, *F*_*(1, 45)*_ = 12.62, *p* = 0.0009; treatment, *F*_*(2, 45)*_ = 15.30, *p* < 0.0001).On the other hand, the proportion of CD68^+^CD206^+^ cells relative to CD206^+^ cells was not significantly different regardless of region and treatment, ranging from 50 to 70 % (two-way ANOVA; interaction, *F*_*(2, 45)*_ = 1.370, *p* = 0.2645; region, *F*_*(1, 45)*_ = 1.174, *p* = 0.2844; treatment, *F*_*(2, 45)*_ = 2.263, *p* =0.1157).

**Table 4.** Proportion of CD68^+^CD206^+^ cells relative to CD68^+^ and CD206^+^ cells as shown in Figure 3g. Data are reported as Mean ± SEM. N = 4, 3, 6 animals with two tissue slices analyzed for untreated TBI, sham, ES.

|  |  | Proportion of CD68<sup>+</sup>CD206<sup>+</sup> (%) |  |  | p-values |  |  |
| --- | --- | --- | --- | --- | --- | --- | --- |
|  |  | TBI | sham | ES | TBI vs. sham | TBI vs. ES | sham vs. ES |
| relative to CD68<sup>+</sup> | ROI<sub>1</sub> | 41.0 ± 11.7 | 44.9 ± 13.4 | 60.0 ± 6.7 | *p* = 0.9448 | *p* = 0.1785 | *p* = 0.3952 |
| | ROI<sub>2</sub> | 11.0 ± 4.6 | 4.3 ± 0.8 | 61.0 ± 5.3 | *p* = 0.8474 | ***p* < 0.0001** | ***p* < 0.0001** |
| relative to CD206<sup>+</sup> | ROI<sub>1</sub> | 54.3 ± 7.9 | 68.8 ± 9.1 | 63.9 ± 4.1 | *p* = 0.4180 | *p* = 0.5956 | *p* = 0.8901 |
| | ROI<sub>2</sub> | 49.3 ± 12.0 | 48.9 ± 9.1 | 68.8 ± 3.7 | *p* = 0.9994 | *p* = 0.1175 | *p* = 0.1545 |
| two-way ANOVA with Tukey's post-hoc analysis |  |  |  |  |  |  |  |

### ES treatment increased abundance of CD206^+^ microglial cells

To specify the type of CD206^+^ immune cells that responded to ES, we performed a flow cytometric analysis on cells obtained from bulk tissue of the entire perilesional cortex and hippocampus (Figure 4 and S3). CD45 antibody was used to differentiate resident microglia (CD45^low^) from blood-derived leukocytes (CD45^high^) (Febinger et al., 2015; Posel et al., 2016).The proportion of CD45^low^ cells to CD45^+^ cells increased significantly after ES (85.5 ± 4.4 %) compared to those in untreated TBI (67.2 ± 7.1 %; p = 0.0230) and sham (66.9 ± 6.0 %; *p* = 0.0138; *p* = 0.9998 for TBI vs. sham) groups (Welch ANOVA test with Dunnett post-hoc; *W*_*(2, 5*.*300)*_ = 12.93, *p* = 0.0092) (Figure 4a), whereas the proportion of CD45^high^ cells decreased in the ES group (Welch ANOVA test with Dunnett post-hoc; *W*_*(2, 5*.*330)*_ = 10.53, *p* = 0.0141) (TBI = 32.2 ± 7.5 %; sham = 32.7 ± 7.0; ES = 14.7 ± 4.5 %; *p* = 0.9993, *p* = 0.0323, and *p* = 0.0239 for TBI vs. sham, TBI vs. ES, and sham vs. ES) (Figure 4b). This may suggest ES induced recruitment of microglia while leukocytes are being suppressed in the perilesional cortex and hippocampus. When we measured CD206^+^ cells within each CD45^low^ or CD45^high^ subpopulation, only CD45^low^ cells showed treatment-dependent differences among phenotypic subpopulations (Welch ANOVA test with Dunnett post-hoc; for Figure 4c, *W*_*(2, 4*.*620)*_ = 220.2,*p* < 0.0001; for Figure 4d, *W*_*(2, 4*.*046)*_ = 5.365, *p* = 0.0728). First, in the untreated TBI group, 43.6 ± 4.0 % of CD45^low^ cells expressed CD206, and the sham group showed a significantly increased CD206^+^ cohort within CD45^low^ cells (52.7 ± 1.4 %) compared to the untreated TBI group (*p* = 0.0016) (Figure 4c). In the ES group, 77.4 ± 1.7 % of CD45^low^ cells were CD206^+^, displaying a statistical significance against the TBI (*p* < 0.0001) and the sham group (*p* < 0.0001). Unlike CD45^low^ microglia, our measurement of the CD206^+^ subpopulation within CD45^high^ leukocytes did not exhibit statistical differences among all three groups (TBI = 53.3 ± 9.9 %; sham = 62.6 ± 3.3 %; ES = 74.0 ± 6.5 %; p = 0.3364, 0.0533, and 0.1532 for TBI vs. sham, TBI vs. ES, and sham vs. ES) (Figure 4d).Lastly, based on the normalization of each population to CD45^+^ cells, it was found that only ES treatment induced significant phenotypic polarization of the observed immune cells, predominantly marked by an increase in CD206^+^ microglia (Welch ANOVA test with Dunnett post-hoc; for CD45^low^CD206^+^/CD45^+^, *W*_*(2, 4*.*785)*_ = 53.21, *p* = 0.0005; for CD45^high^CD206^+^/CD45^+^, *W*_*(2, 3*.*611)*_ = 4.814, *p* = 0.0958) (CD45^low^CD206^+^/CD45^+^ for TBI = 29.5 ± 5.6 %, sham = 35.2 ± 3.6 %, and ES = 66.2 ± 4.5 %;*p* = 0.1424, *p* = 0.0007, and *p* = 0.0016 for TBI vs. sham, TBI vs. ES, and sham vs. ES) (Figure 4e).

**FIGURE 4.**
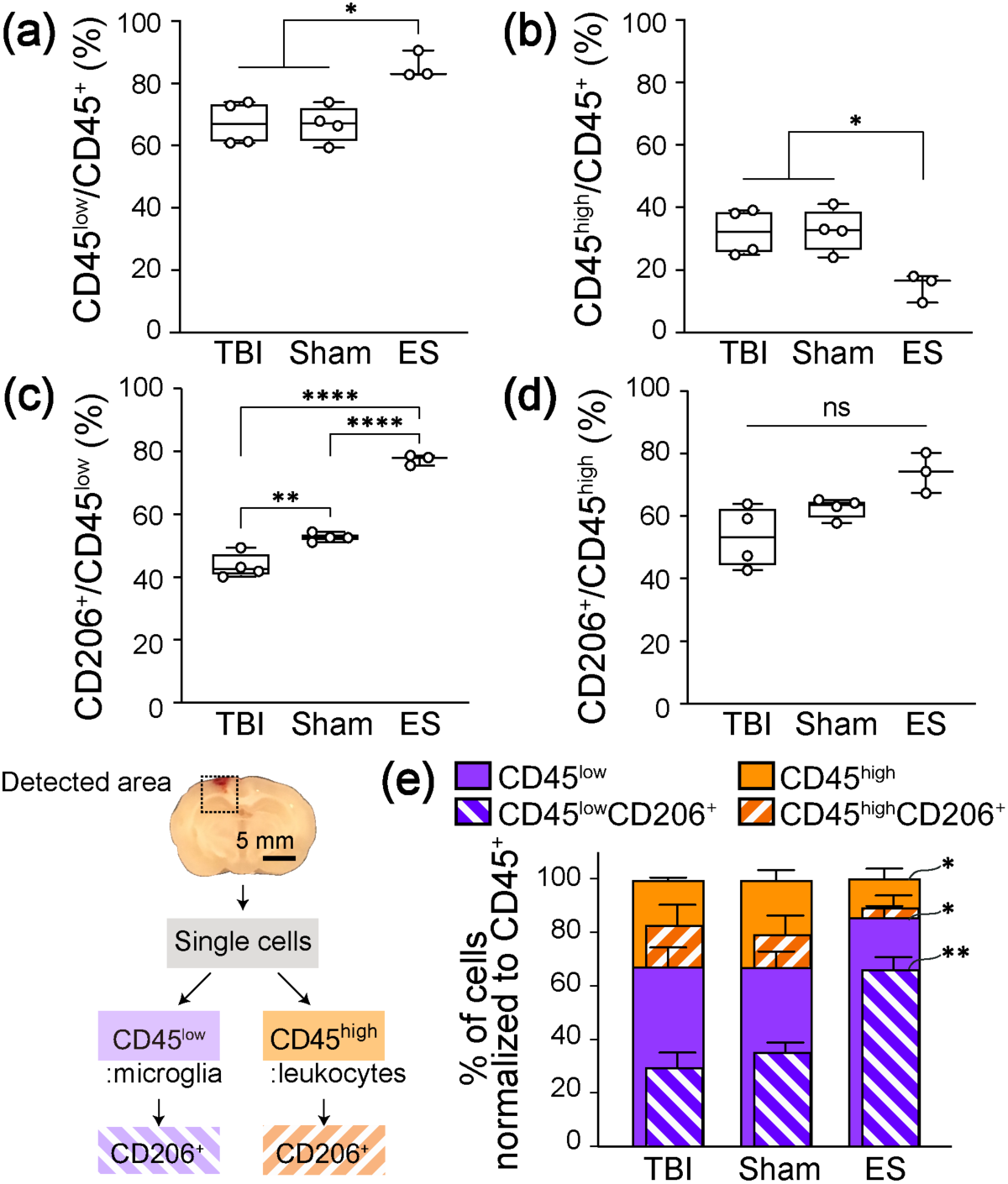
Flow cytometric analysis to identify the population of immune cell subtypes and their CD206^+^ population at 7 days post-TBI. The detected area for flow cytometry included the perilesional cortex and the ipsilateral hippocampus. The sliced tissue shows the coronal plane of the center of the injury. (a, b) Following the ES treatment, the change in the proportion of immune cells that (a) CD45^low^ microglia increase, while the (b) CD45^high^ leukocytes decrease. (c, d) The proportion of CD206^+^ cells to the CD45^low^ and CD45^high^ cells, respectively. In sham and ES groups, more microglia expressed CD206 (c), whereas leukocytes did not show significant phenotypic changes (d). (e) The normalized graph of immune cell subtypes and their CD206^+^ population to the CD45^+^ cells. ES treatment showed a significant increase in the CD45^low^CD206^+^ cells. Graphs are shown as Min to Max with all points for Figure a-d and mean ± SD for Figure e. N = 4, 4, 3 animals analyzed for untreated TBI, sham, ES; * *p* < 0.05, ** *p* < 0.01, *** *p* < 0.001, **** *p* < 0.0001, Welch ANOVA tests with Dunnett post-hoc analysis.

### ES treatment increased the populations of Nestin^+^ and Nestin^+^GFAP^+^ cells in the ipsilateral cortex

To investigate the endogenous healing response in the perilesional cortex by ES after TBI, immunohistochemistry was performed on our collected tissues to measure the expression levels of Nestin and glial fibrillary acidic protein (GFAP), a general marker for NSCs and astrocytes, respectively (Figure 5a). Regarding regional distribution, GFAP^+^ cells were observed throughout the entire cortex in all experimental groups. Few Nestin^+^ cells were found overall, located primarily along the perilesional rims in the untreated TBI and sham groups. In contrast, in the ES group, Nestin^+^ cells were spread liberally from the perilesional rims to the distant cerebral cortex. Moreover, some fraction of Nestin^+^ cells co-expressed GFAP. Using confocal z-stack colocalization analysis in coronal planes, we confirmed that the majority of the Nestin^+^ voxels were colocalized with GFAP in all experimental groups (Figure 5b). In the TBI and sham groups, in particular, not all GFAP^+^ voxels were positive for Nestin, but in the ES group, GFAP^+^ voxels mostly coincided with Nestin expression.

**FIGURE 5.**
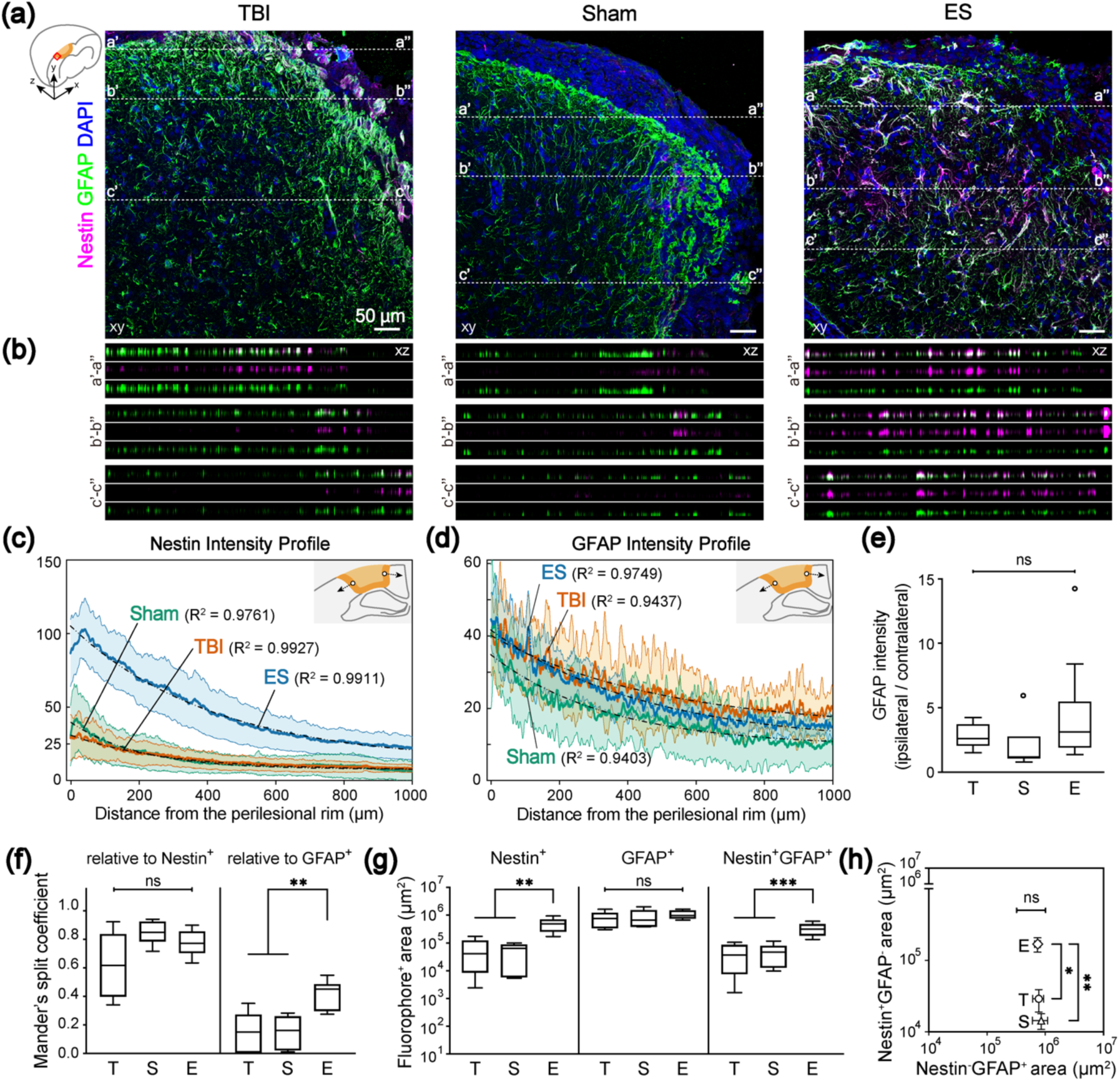
Increased Nestin^+^ and Nestin^+^GFAP^+^ cells in the perilesional cortex at 7 days post-TBI by ES treatment. (a) Representative z-projection (xy) confocal images of the perilesional cortex. Magenta, green, and blue represent Nestin, GFAP, and DAPI, respectively. In Figure a, three representative positions (a-a’, b-b’, c-c’) were chosen to show the differential voxel distribution and their orthogonal views (xz) of the dotted lines in each z-projection image are in Figure b. (b) Images are listed in order of merged, magenta, and green from the top. From the xz-section images, it was confirmed that Nestin^+^ voxels are coincident with GFAP^+^ voxels (The total depth of z-stack images: 4, 4, and 7 *μm* in TBI, sham, and ES groups. The step size: 1 *μm*). Mean profiles of (c) Nestin and (d) GFAP intensity and 95 % confidence intervals (shaded bands). The schematics for the analysis of gradual changes in Nestin and GFAP expression are illustrated in each figure: the blank circle and the arrow indicate the initial point and the direction of analysis, respectively. In all experimental groups, mean intensity was fit to the one-phase exponential decay curve, and their R-squared values are shown on the graph. (e-g) T, S, and E on the x-axis represent untreated TBI, sham and ES groups. (e) GFAP intensity ratio of ipsilateral/contralateral cortex was larger than 1, but was not significant among groups, indicating that TBI induced astrocyte activation at the perilesional cortex, but sham or ES operations did not induce additional activation. (f) The degree of GFAP- and Nestin-concurrent signals with Nestin (left) and GFAP (right), respectively. There was no significant difference in the Mander’s split coefficient relative to Nestin^+^ pixels, whereas more GFAP^+^ cells appeared to express Nestin signals in the ES group compared to the TBI and sham groups. (g) Quantitative analysis of Nestin^+^, GFAP^+^, Nestin^+^GFAP^+^ area. Nestin^+^ and Nestin^+^GFAP^+^ area showed a treatment-dependent change, an increase by the ES treatment. (h) Nestin^-^GFAP^+^ area (x-axis) and Nestin^+^GFAP^-^ area (y-axis). T, S, and E next in graph represent untreated TBI, sham, and ES groups. There was no significant difference in Nestin^-^ GFAP^+^ area, whereas Nestin^+^GFAP^-^ area of ES group was significantly larger than untreated TBI and sham groups. Data are represented as mean ± SD for Figure e-g, and mean ± SEM for Figure h. N = 4, 3, 6 animals with two tissue slices analyzed for untreated TBI, sham, ES; * *p* < 0.05, ** *p* < 0.01, *** *p* < 0.001, Kruskal-Wallis one-way ANOVA with Dunn’s post-hoc analysis for Figure e and Welch ANOVA with Dunnett post-hoc analysis for others.

Next, the spatial distribution of Nestin (Figure 5c) and GFAP (Figure 5d) intensity profiles from the perilesional rim further into the cortex was identified. First, in both the untreated TBI and sham groups, Nestin was predominantly expressed in the perilesional rims, but not in areas distant from perilesional rim, whereas in the ES group, a substantial increase in Nestin intensity in the perilesional cortex was observed with a gradual decrease away from the injury site (Figure 5c). We also confirmed that the GFAP intensity was at its maximum level near the perilesional rim and decreased exponentially (Figure 5d). The mean Nestin or GFAP intensity profiles were best fit to the one-phase exponential decay curve (Figure S4 and Figure 5c,d). From the fitting curves of Nestin and GFAP intensity profiles, two values were obtained: rate constant (*k*) that measures the rate of intensity-decrease over a distance, and half-life (*in* (2)/*k*) that represents a distance required for intensity to reduce to half of its initial value (Figure S4c). In Nestin fitting curve, the fitted values of rate constant (TBI = 0.0035/μm, sham = 0.0049/μm, ES = 0.0024/μm) and half-life (TBI = 196.8 μm, sham = 141.7 μm, ES = 289.9 μm) demonstrated that Nestin intensity rapidly decreased in sham, TBI, and ES, sequentially. In the curve fit for GFAP, as compared to the Nestin-fitting results, the rate constants of GFAP in all experimental groups (TBI = 0.0020/μm, sham = 0.0029/μm, ES = 0.0022/μm) were observed to be reduced and the half-life of the GFAP-fitting curve (TBI = 342.1 μm, sham = 237.9 μm, ES = 318.3 μm) increased, indicating that GFAP intensity persisted further than Nestin intensity. In addition, to determine whether the injury and/or treatment induced an astrocytic activation, GFAP expression in the ipsilateral cortex compared to that of the contralateral side was analyzed (Figure 5e). The relative expression of GFAP in the ipsilateral cortex appeared to increase on average, but there were no significant differences between all three groups (*p* = 0.0503, *Kruskal-Wallis statistic* = 5.981, *df* = 2; TBI = 2.81 ± 1.02; sham = 2.00 ± 1.99; ES = 4.45 ± 4.03; *p* = 0.1862, *p* > 0.9999, and *p* = 0.0516 for TBI vs. sham, TBI vs. ES, and sham vs. ES).

Furthermore, within 0-2 mm from the perilesional rims, we used Mander’s split coefficients (Zinchuk et al., 2007) to analyze the degree of GFAP and Nestin colocalization, which corresponds to the proportion of colocalized signal relative to overall fluorophore^+^ pixels (Figure 5f).The colocalization coefficient relative to Nestin^+^ pixels appeared not significantly different in all treatment conditions (Welch ANOVA test with Dunnett post-hoc; *W*_*(2, 11*.*70)*_ = 3.716, *p* = 0.0563; TBI = 0.619 ± 0.225, sham = 0.848 ± 0.082, ES = 0.777 ± 0.083; *p* = 0.0721,*p* = *0*.2361, and *p* = 0.3075 for TBI vs.sham,TBI vs. ES, and sham vs. ES), whereas the colocalization coefficient relative to GFAP^+^ pixels increased significantly in the ES group (Welch ANOVA test with Dunnett post-hoc; *W*_*(2, 11*.*97)*_ = 15.55, *p* = 0.0005; TBI = 0.148 ± 0.133, sham = 0.149 ± 0.112, ES = 0.410 ± 0.100; *p* > 0.9999, *p* = 0.0014, and *p* = 0.0025 for TBI vs. sham, TBI vs. ES, and sham vs. ES).

Next,we measured the (projected) areas where Nestin or GFAP signals are observed within the same cortical ROI used for the above colocalization analysis (Figure 5g).Based on the detected Nestin fluorescent signal, the significantly increased population of Nestin^+^ cells in the ES group was confirmed (498,738 ± 263,474 μm^2^), when compared to those of the untreated TBI (64,462 ± 62,754 μm^2^; *p* = 0.0015) and sham rats (53,664 ± 39,387 μm^2^; *p* = 0.0011) (TBI vs. sham: p = 0.9700) (Welch ANOVA test with Dunnett post-hoc; *W*_*(2, 13*.*69)*_ = 13.13, *p* = 0.0007). Comparatively, the GFAP^+^ area was larger than the Nestin^+^ area, displaying no statistical significance between groups (Welch ANOVA test with Dunnett post-hoc; *W*_*(2, 10*.*58)*_ = 0.7981, *p* = 0.4755; TBI = 820,341 ± 476,882 μm^2^, sham = 902,247 ± 637,979 μm^2^, ES = 1,071,960 ± 342,906 μm^2^; *p* = 0.9904, *p* = 0.5289, and *p* = 0.9043 for TBI vs. sham, TBI vs.

ES, and sham vs. ES). Furthermore, compared to untreated TBI and sham groups, a significant increase in the regions with both Nestin and GFAP signals was confirmed in the ES group (Welch ANOVA test with Dunnett post-hoc; *W*_*(2, 13*.*27)*_ = 15.23, *p* = 0.0004; TBI = 44,166 ± 38,756 μm^2^,sham = 49,662 ± 38,907 μm^2^, ES = 330,314 ± 155,412 μm^2^; *p* = 0.9907, *p* =0.0007, and *p* = 0.0006 for TBI vs. sham, TBI vs. ES, and sham vs. ES). Next, the areas of Nestin^+^GFAP^-^ and Nestin^-^GFAP^+^ regions were calculated by subtracting Nestin^+^GFAP^+^ area from Nestin^+^ and GFAP^+^ area, respectively (Figure 5h).While the Nestin^+^GFAP^-^ area in the ES group was significantly larger than that of untreated TBI and sham groups (Welch ANOVA test with Dunnett post-hoc; *W*_*(2, 11*.*86)*_ = 8.354, *p* = 0.0054; TBI = 28,935 ± 28,389 μm^2^, sham = 14,348 ± 8,799 μm^2^, ES = 168,424 ± 120,881 μm^2^; *p* = 0.4702, *p* = 0.0156, and *p* = 0.0087 for TBI vs. sham, TBI vs. ES, and sham vs. ES), whereas there were no significant differences in Nestin^-^GFAP^+^ area across the groups (Welch ANOVA test with Dunnett post-hoc; *W*_*(2, 9*.*943)*_ = 0.08872, *p* = 0.9158; TBI = 776,176 ± 477,085 μm^2^, sham = 852,585 ± 613,177 μm^2^, ES = 741,646 ± 276,558 μm^2^; *p* = 0.9915, *p* = 0.9968, and *p* = 0.9633 for TBI vs. sham, TBI vs. ES, and sham vs. ES).

### Hippocampal Nestin^+^ cells exhibited distinct morphologies from those in the ipsilateral cortex

Next, we analyzed Nestin and GFAP expression in the hippocampal region (Figure 6). An increased number of Nestin^+^ cells in the ES group was observed also in the hippocampal region. However, unlike Nestin^+^ cells in the ipsilateral cortex, the hippocampal Nestin^+^ cells were not predominantly colocalized with GFAP proteins. Here, three marked characteristics of hippocampal Nestin^+^ cells were identified based on the GFAP expression and cellular morphology. Many Nestin^+^GFAP^-^ cells displayed vessel-like morphology that featured plump cell bodies, showing 2-3 cells aligned as if they were adjoined to one another (Figure 6b). In addition, long and slender processes of GFAP^+^ cells appeared to be in contact with the vessel-like Nestin^+^GFAP^-^ cells, as if the end-feet of astrocytes were wrapping around the blood vessels. Such vessel-like Nestin^+^GFAP^-^ cells were shown in all experimental groups. Nestin^+^ cells that featured multiple processes were denoted as fibrous Nestin^+^GFAP^-^ cells (Figure 6c) or fibrous Nestin^+^GFAP^+^ cells (Figure 6d).

**FIGURE 6.**
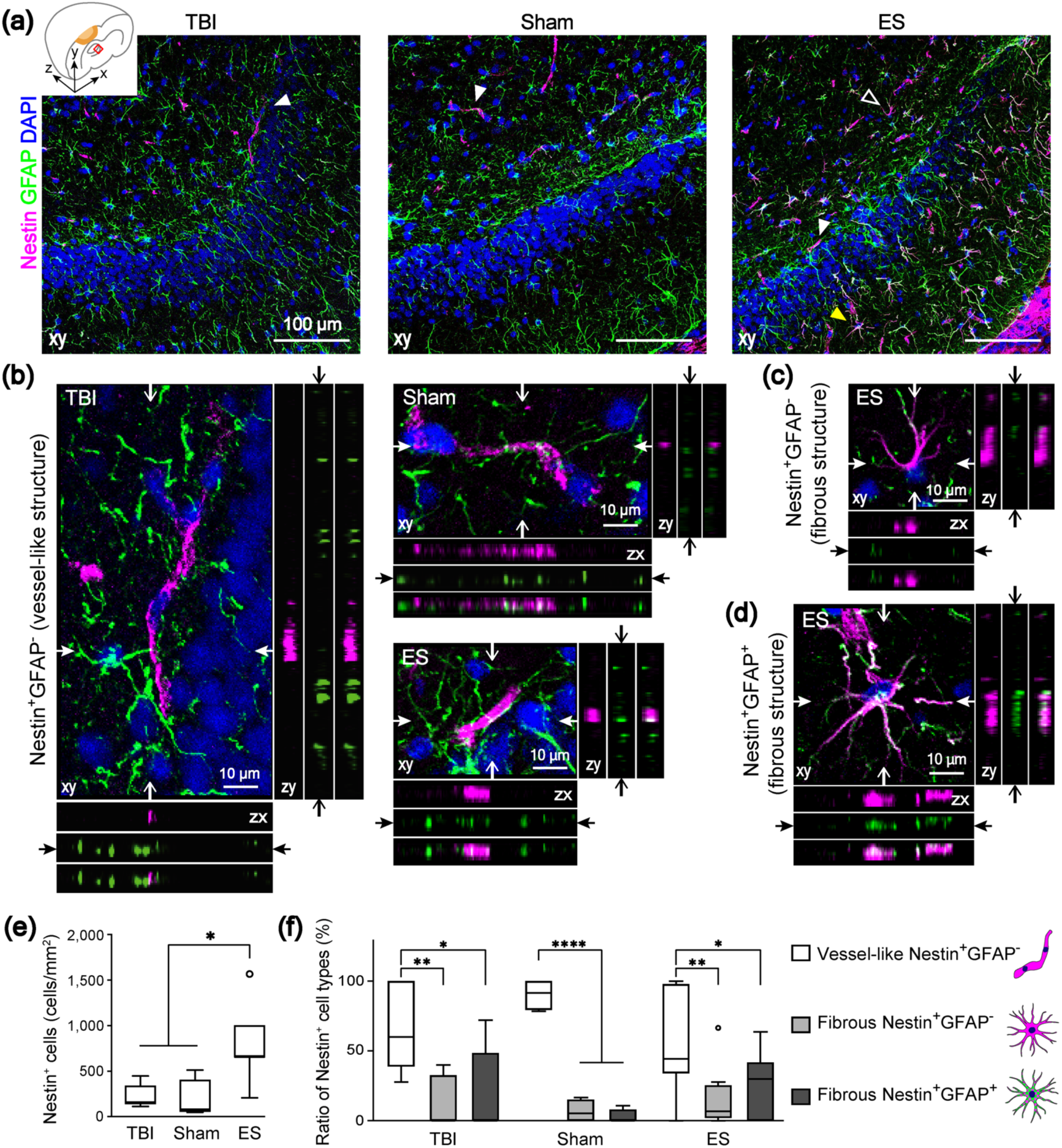
The phenotypes of Nestin^+^ cells in the ipsilateral hippocampus at 7 days post-TBI. Magenta, green, and blue represent Nestin, GFAP, and DAPI, respectively. (a) Representative z-projection (xy) confocal images of the ipsilateral hippocampus. (b-d) Three representative phenotypes of Nestin^+^ cells, marked with arrowheads in Figure a. In each z-projection (xy) image, the orthogonal views of the position indicated by arrows are on the bottom (zx) and right (zy) panels. In each bottom and right panels, images are listed in order of magenta, green and merged from the top and left, respectively (The total depth of z-stack images: 4 *μm*, the step size: 1 *μm*). (b) White arrowhead in Figure a indicates Nestin^+^GFAP^-^ cells, featuring vessel-like structure. A white empty or yellow arrowhead in Figure a indicates fibrous Nestin^+^GFAP^-^ cells (c) or fibrous Nestin^+^GFAP^+^ cells (d) that feature multiple processes. (e) Quantitative analysis of the number of Nestin^+^ cells in the ROI of a hippocampus revealed that ES treatment increased the number of Nestin^+^ cells. (f) The phenotypic ratio of Nestin^+^ cells within a group. Nestin^+^ cells found in the hippocampus were mostly vessel-like structural Nestin^+^GFAP^-^ cell. Data are represented as mean ± SD. N = 4, 3, 6 animals with two tissue slices analyzed for untreated TBI, sham, ES; * *p* < 0.05, ** *p* < 0.01, *** *p* < 0.001, **** *p* < 0.0001, Kruskal-Wallis one-way ANOVA with Dunn’s post-hoc analysis in Figure e, and Two-way ANOVA tests with Tukey post-hoc analysis in Figure f.

We quantified the number of Nestin^+^ cells per field in the dentate gyrus granule cell layer, and found that the ES treated group had a high number of Nestin^+^ cells in the ipsilateral hippocampal region (*p* = 0.0016, *Kruskal-Wallis statistic* = 9.737, *df* = 2), displaying a statistical significance against sham but not against TBI (TBI = 224.0 ± 133.4 cells/mm^2^,sham = 180.0 ± 222.2 cells/mm^2^, ES = 800.0 ± 417.9 cells/mm^2^; p > 0.9999, = 0.0789, = 0.0118 for TBI vs. sham, TBI vs. ES, sham vs. ES) (Figure 6e). Then, the relative ratios of the three structural sub-phenotypes were quantified (two-way ANOVA with Tukey’s post-hoc; interaction, *F*_*(4, 45)*_ = 1.980, *p* = 0.1138; sub-phenotypes, *F*_*(2, 45)*_ = 28.87, *p* < 0.0001; treatment, *F*_*(2, 45)*_ = 6.01e-009, *p* > 0.9999) (Figure 6f). All experimental groups showed a significantly higher proportion of Nestin^+^GFAP^-^ vessel-like structural cells among all Nestin^+^ cells (TBI =67.6 ± 31.8 %, sham = 90.4 ± 11.2 %, and ES = 57.2 ± 35.4 %). Relatively, the proportion of fibrous Nestin^+^GFAP^-^ cells among all Nestin^+^ cells (TBI = 13.0 ± 18.6 %, sham = 6.9 ± 8.3 %, and ES = 16.4 ± 21.3 %) and fibrous Nestin^+^GFAP^+^ cells (TBI = 19.4 ± 31.4 %, sham = 2.7 ± 5.4 %, and ES = 26.5 ± 22.5 %) significantly decreased (TBI: *p* = 0.0032, *p* = 0.0101, and *p* = 0.9115; sham: *p* < 0.0001, *p* < 0.0001, and *p* = 0.9694; ES: *p* = 0.0031, *p* = 0.0313, and *p* = 0.6652; *p*-values in each group represent vessel-like cells vs.fibrous Nestin^+^GFAP^-^ cells, vessel-like cells vs. fibrous Nestin^+^GFAP^+^ cells, and fibrous Nestin^+^GFAP^-^ cells vs. fibrous Nestin^+^GFAP^+^ cells, respectively).

### Neuronal viability in the perilesional cortex increased in groups with an implanted electrode (sham & ES) following TBI

To assess the effect of ES on the primary trauma to neural tissues after TBI, the presence of NeuN, a marker for mature neurons, was analyzed by immunohistochemistry (Figure 7a). Cells in the core of the cortical lesion exhibited very low NeuN immunoreactivity, indicative of an increase in injured neurons. To quantify the progress of neuronal loss near the perilesional cortex, the number of NeuN^+^ cells within 250 *μm* from the injury in ROI containing at least 100 nuclei was measured (Figure 7b). In all experimental groups, the number of NeuN^+^ cells decreased in the perilesional cortex compared to its uninjured contralateral side, implying that TBI gave rise to neuronal loss, but the level of discrepancy was treatment dependent (*p* = 0.0064, *Kruskal-Wallis statistic* = 9.024, *df* = 2). In untreated TBI group, 54.3 ± 13.9 % of NeuN^+^ cells relative to the contralateral side was observed. In the perilesional cortex of sham and ES, significantly increased NeuN^+^ expression was confirmed compared to the untreated TBI group (sham = 75.5 ± 9.8 %, ES = 77.7 ± 20.5 %; *p* = 0.0461 for TBI vs. sham, *p* = 0.0211 for TBI vs. ES), but there were no significant differences between the sham and ES groups (*p* > 0.0999). We also quantified the progress of neuronal loss in the CA3 region close to the dentate gyrus of hippocampus where NSCs are believed to initiate endogenous regeneration following the injury. In all experimental groups, the NeuN^+^ cell population in the ipsilateral CA3 was lower than that in the contralateral side, of which the relative fractions were 81.9 ± 39.3 % of for TBI, 88.1 ± 25.7 % for sham, and 92.8 ± 18.4 % for ES with no statistical significance among all groups (Welch ANOVA test with Dunnett post-hoc; *W*_*(2, 13*.*99)*_ = 0.3721, *p* = 0.6959).

**FIGURE 7.**
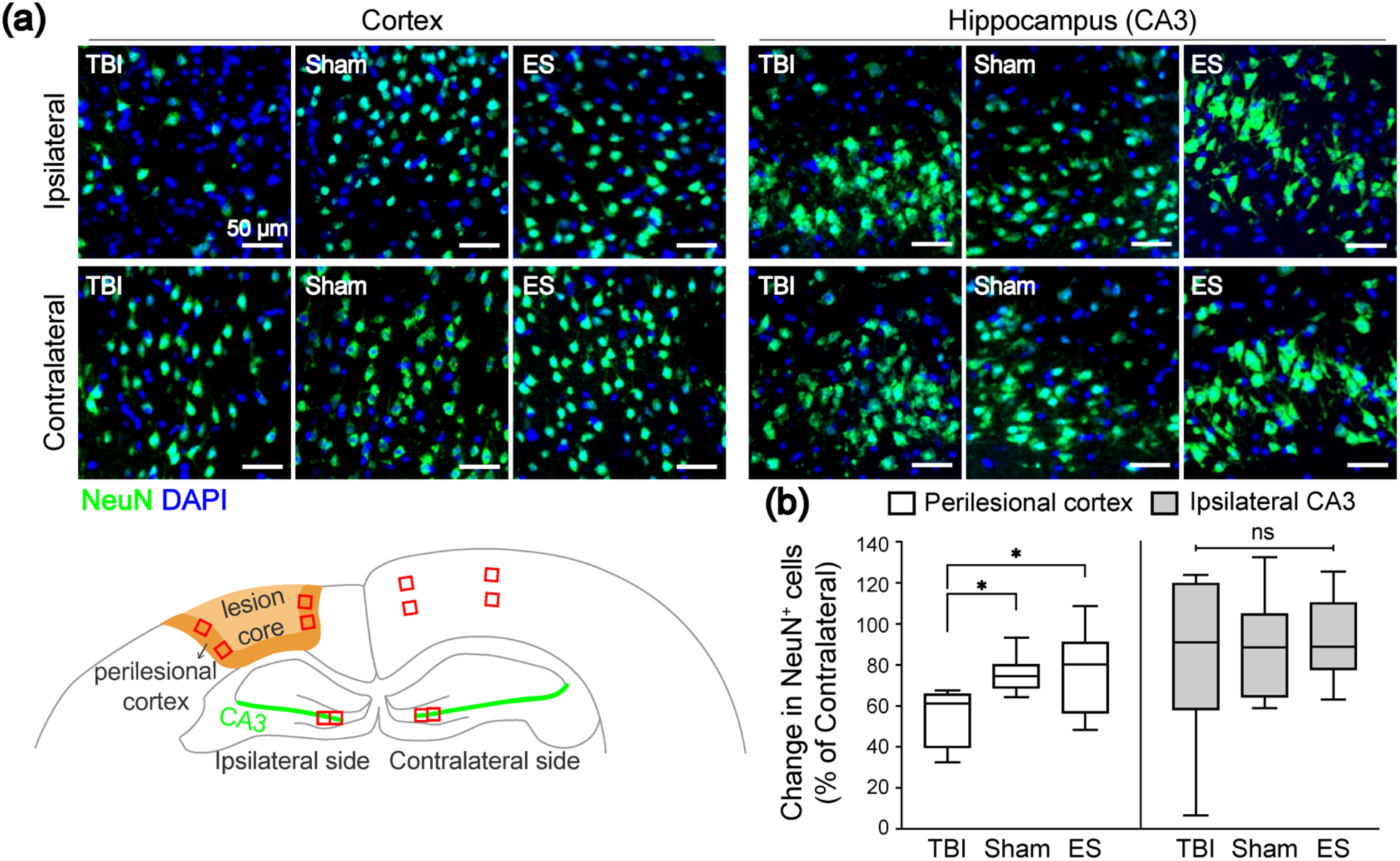
Effect of treatment on neuronal viability in the ipsilateral cortex and hippocampus at seven days post-TBI. (a) Representative fluorescence images of NeuN^+^ neurons (green) in the cortex (left three columns) and CA3 in the hippocampus (right three columns). The images in the first row and second row represent the NeuN in the ipsilateral and contralateral sides, respectively. (b) Quantitative analysis of the change in the number of NeuN^+^ cells in the ipsilateral ROI compared to the contralateral side. In the ipsilateral cortex, NeuN^+^ cells remained more in the sham and ES groups compared to untreated TBI group. On the other hand, the changes of NeuN^+^ cells in the ipsilateral CA3 region were not significantly different across the groups. For the analysis of NeuN^+^ cells, four ROIs were taken from each ipsilateral and contralateral cortices, and two ROIs were taken from each ipsilateral and contralateral CA3 regions. N = 4, 3, 6 animals with two coronal sections analyzed for untreated TBI, sham, ES; * *p* < 0.05, Kruskal-Wallis one-way ANOVA with Dunn’s post-hoc analysis.

## DISCUSSION

Neuronal damage and death are the most critical pathological consequences in TBI, incurring long-lasting and functional impairment in the brain. For the improved recovery of brain function, neuronal death should be minimized as well as secondary, deleterious effects by shifting the injury-associated cellular and inflammatory milieu. In this study, we showed the potential use of an invasive ES as a means to therapeutically improve this milieu following TBI. In this study, an increase in CD206^+^ cells was observed after ES in the perilesional cortex, and the distribution of these cells shifted deeper into the cortex than in control and sham animals. ES was also observed to increase the relative abundance of CD206^+^ microglial cells, and the populations of Nestin^+^ and Nestin^+^GFAP^+^ cells in the injury-ipsilateral cortex. The Nestin^+^ cells in the hippocampus also exhibited distinct morphologies from those in the ipsilateral cortex after ES. Lastly, after TBI, neuronal viability in the perilesional cortex was observed to increase in animals with an implanted electrode regardless of ES condition. Please note, conclusions in this study are limited to male rats as sex differences were not studied.

We chose to use invasive electrodes to localize the effects of the ES to the injury site. To minimize any side effects during electric stimulation, a balanced-biphasic stimulation mode was used such that accumulation of charges and toxic byproducts would be minimized around the electrode (Merrill et al., 2005). We also considered the timing of ES treatment. Immunomodulation in the early stages of brain damage has emerged as one of the important therapeutic targets because phenotypic alterations during acute neuroinflammation (Kumar et al., 2016; Wang et al., 2013) can critically attribute to chronic pathologies, including the induction of neuronal death or attenuation of the neurogenesis (Loane, Kumar, Stoica, Cabatbat, & Faden, 2014; Schimmel, Acosta, & Lozano, 2017). The positive impact of appropriate timing of immunomodulation, especially early in the regeneration process, has been shown in multiple peripheral nerve studies (Mokarram et al., 2017; Mukhatyar et al., 2014; Ydens et al., 2012). Additionally, NSCs have been shown to be involved in the early healing process by being activated within three days following TBI (Itoh et al., 2005; Yi et al., 2013). Based on these observations, we decided to apply the ES on the 2nd day post-TBI to modulate the phenotypes of both immune cells and NSCs.

Regarding neuroinflammation, we first assessed the M2 phenotype population by counting cells that expressed CD68 and/or CD206, which revealed differential spatial distribution of phenotypic markers that depended on cortical depth relative to injury lesion (Figure 2 and Figure 3). Following TBI, CD68^+^ cells were shown to be distributed throughout the entire perilesional cortex and corpus callosum excluding the hippocampus, whereas CD206^+^ cells were mostly observed near the cortical surface. Perego, et al., showed differential spatial patterns of immune cell markers and M2 phenotype markers in ischemic stroke, similar to our results (Perego, Fumagalli, & De Simoni, 2011). Moreover, they observed phagocytosed neurons by CD11b/CD68 double positive immune cells in the region where the M2 phenotype cells were not found. Considering that CD68 is not only an immune cell marker but also used as a phagocytic cell marker, these collective data suggest that innate healing processes might be limited to the cortical surface under natural circumstances after brain damage. In our study, we found that ES treatment could increase CD206 expression regardless of CD68 expression in the region of 1-2 mm from the cortical surface to levels as high as was found in the region of 0-1 mm from there (Figure 3d-f)–and without a change in the density and spatial distribution of CD68^+^ cells, whereas in the sham operation this was not observed (Figure 3c).

However, in our flow cytometric analysis of CD45^+^ cells (leukocytes and microglia) from the entire perilesional cortex and hippocampus, the presence of an implanted electrode was sufficient to increase the proportion of CD206^+^ microglia relative to untreated TBI group (Figure 4), though this was not corroborated using CD68 immunohistochemistry (Figure 2,3). Even so, the level of CD206 expression in microglia grew markedly with ES compared to untreated TBI and sham groups (Figure 4c), whereas leukocytes did not respond with an M2-like phenotype shift even after electrical stimuli (Figure 4d). Based on these results, we summarized the phenotypic marker expression (Figure 8a-c). We found that the sham operation slightly increased CD68^+^CD206^+^ cells in the region of 0-1 mm from the cortical surface compared to untreated TBI on average, though there was no significance (Table 3 and Figure 3e). We assume that this slight change was due to microglial response, which was discriminable using flow cytometry (Figure 8b). A remarkable increase of CD206 expression in the CD68^+^ (Figure 3e) and CD68^-^ (Figure 3f) populations after ES was shown in Figure 8c, which is thought be driven by phenotypic alterations in CD45^low^ microglia as identified with flow cytometry (Figure 4c).

**FIGURE 8.**
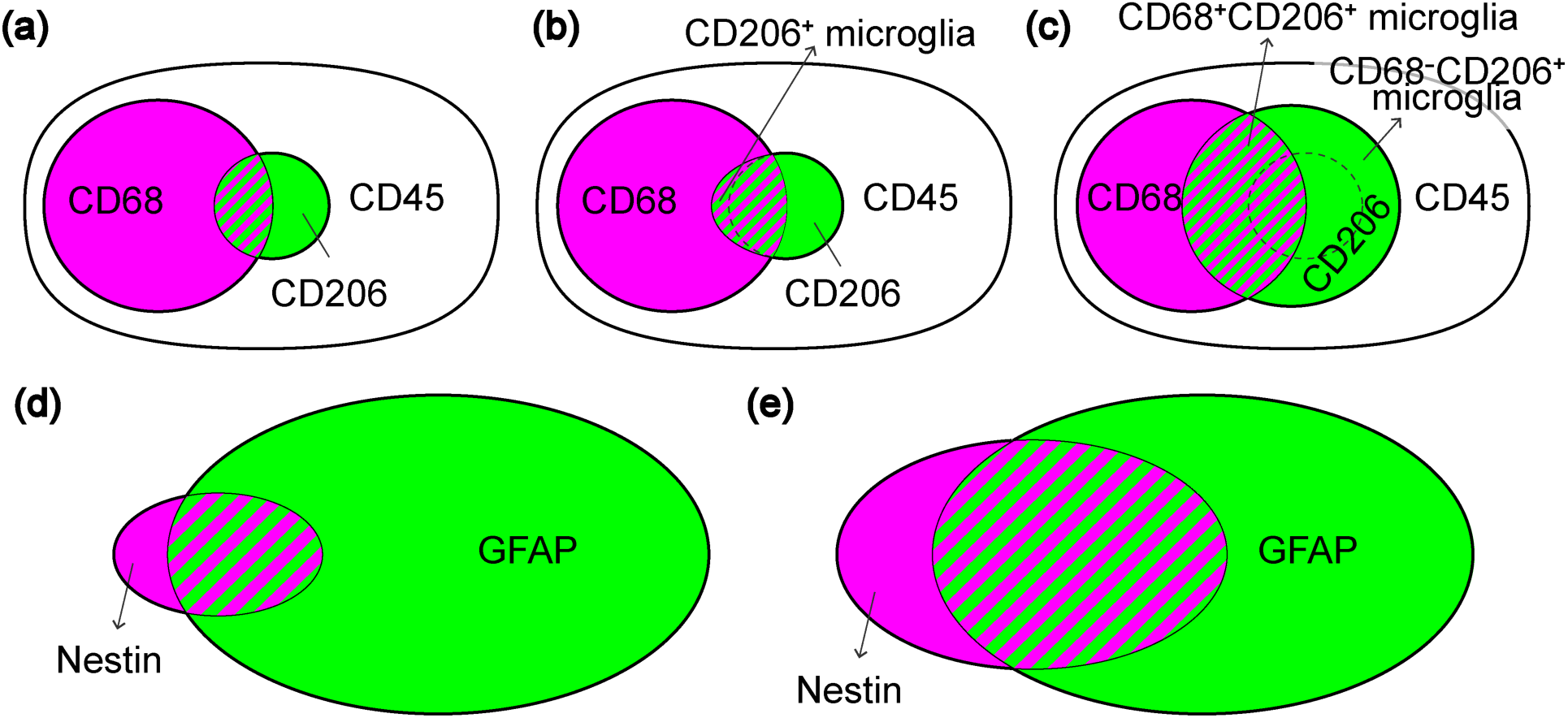
Summary of inflammatory marker and Nestin/GFAP expression at seven days post-TBI. (a-c) CD45, CD68, and CD206 expression identified by immunohistochemistry (Figure 2,3) and flow cytometry (Figure 4). Shaded bands represent CD68^+^CD206^+^ population among CD45^+^ cells. CD45 is a general marker for microglia and leukocytes. CD68 is expressed in phagocytic microglia and leukocyte subtypes (macrophage/monocyte). (a) The Venn-diagram shows a small CD206^+^ population among either CD68^+^ or CD45^+^ cells in untreated TBI. (b) In the sham group, increased CD206^+^ microglia confirmed by flow cytometry is depicted within CD68 population, given the slight increase in CD68^+^CD206^+^ population in ROI_1_ (Figure 3e). (c) By ES treatment, CD206 population greatly increases among both CD68^+^ and CD68^-^ cells compared to untreated TBI and sham groups, which is shown to be induced by microglia. (d, e) Nestin and GFAP expression. Shaded bands represent Nestin^+^GFAP^+^ population. (d) The Venn-diagram shows smaller Nestin^+^ cells than GFAP^+^ cells in TBI and sham groups (Figure 5g). Also, Nestin^+^GFAP^+^ cells have been derived from either Nestin^+^ and GFAP^+^ cells. (e) In the ES group, Nestin^+^ cells increase compared to untreated TBI and sham groups. The fraction of GFAP^+^ cells among Nestin^+^ cells has not been changed, whereas more Nestin-expressing cells have been observed within GFAP^+^ cells, thereby resulting in more Nestin^+^GFAP^+^ cells (and consequently fewer Nestin^-^GFAP^+^ cells) compared to untreated TBI and sham groups (Figure 5g,h).

The differential response of microglia and leukocytes to ES has yet to be thoroughly explored. Nevertheless, the role of infiltrating leukocytes seems less critical, given that they exist in the brain only temporarily, while microglia are brain-resident immune cells that continually participate throughout all pathological processes in TBI (Donat, Scott, Gentleman, & Sastre, 2017). Therefore, microglia that could have some therapeutic responsiveness to an exogenous electrical stimulus are thought to be a promising cellular target for immunomodulation by ES across both acute and chronic stages of TBI.

We also observed a noticeable increase of cortical Nestin expression after ES (Figure 5). Nestin is known as a marker for NSCs, which are considered to be key players in promoting the regeneration and restoration of the injury site in TBI (Bond et al., 2015). In adult rat TBI, several studies have shown temporal patterns of cortical Nestin expression where Nestin^+^ cells migrate from the neurogenic niche following injury onset to the perilesional cortex, peaking at three days post-TBI and then decreasing (Itoh et al., 2005; Yi et al., 2013). Thus, diminished neural stem cell trafficking in the adult brain is believed to contribute to delayed or incomplete healing.In our analyses of the tissues taken 7d post-TBI, Nestin^+^ cells were rarely observed in untreated TBI and sham rats, whereas a large number of Nestin^+^ cells appeared in the perilesional cortex after ES (Figure 5g left), supporting the notion that ES treatment promoted the maintenance and/or the recruitment of the Nestin^+^ cells from the neurogenic niche.

Interestingly, we confirmed the ES-induced increase in the number of Nestin^+^GFAP^+^ cells in the perilesional cortex (Figure 5g right). We suggest two possibilities for how Nestin^+^GFAP^+^ cells may have been increased by ES. First, Nestin^+^GFAP^+^ cells in our study might have resulted from differentiated Nestin^+^ cells after the brain injury, co-expressing two markers at the same time. However, based on no significance in Mander’s split coefficient relative to Nestin amongst three groups (Figure 5f), we concluded that ES treatment was not likely involved in the process of expressing GFAP in Nestin^+^ cells. The other possibility is GFAP^+^ cells expressing Nestin. Several studies have shown that GFAP^+^ cortical astrocytes could express Nestin after brain injury, after acquiring multipotency (Gabel et al., 2016; Shimada, LeComte, Granger, Quinlan, & Spees, 2012). In the perilesional cortex of our ‘samples, the average GFAP-intensity values (Figure 5e) and GFAP^+^ area (Figure 5g middle) were similar between all three groups, but the Mander’s split coefficient relative to GFAP was noticeably increased by ES. These results indicated that ES treatment may have induced astrocytes to express Nestin while having no immediate impact on the level of GFAP expression. Taken these results together, we suggest that ES treatment facilitates the maintenance/recruitment of NSCs to the cortical injury and the acquisition of stemness in cortical astrocytes, simultaneously (Figure 8d,e).

ES treatment also increased hippocampal Nestin^+^ cells (Figure 6). However, based on phenotypic differences, these Nestin^+^ cells are thought to be different from those in the cortical region. Unlike in the cortical region, the majority of hippocampal Nestin^+^ cells displayed vessel-like phenotypes: 2 or 3 thick cells aligned and adjoined together (Figure 6b). The rest of hippocampal Nestin^+^ cells, with or without GFAP, showed multiple processes, displaying fibrous morphologies distinct from the vessel-like ones (Figure 6c,d). These cells are positively identified as either hippocampal NSCs or those derived from astrocytes, similar to the cortical Nestin^+^ cells. The proportion of these morphologically distinct groups of Nestin^+^ cells was similar in all experimental groups (Figure 6f), suggesting that ES treatment following TBI promoted overall Nestin^+^ cells, with no specificity in any particular morphology of Nestin^+^ cells. While the identity of vessel-like Nestin^+^ cells remains unknown, some recent studies have provided some critical clues as to what they might be. Nakagomi, et al., demonstrated that brain vascular pericytes acquired stem cell-like properties following cortical ischemia (Nakagomi et al., 2015), and that those brain vascular pericytes exhibited similar morphological traits to our vessel-like Nestin^+^ cells that occurred during the recovery following TBI. Based on the morphological similarities, one possibility could be that these vessel-like Nestin^+^ cells were indeed hippocampal pericytes expressing Nestin. In addition, Nestin^+^ pericytes have been previously observed to differentiate into microglia (Sakuma et al., 2016). Thus, our observed increase of vessel-like Nestin^+^ cells by ES (Figure 6e,f) may have contributed to the observed increase in microglial population (Figure 4a), thereby decreasing the relative proportion of leukocytes (Figure 4b).

Overall, the increase in both cortical and hippocampal Nestin^+^ cells by ES can be interpreted in connection with concurrently promoted anti-inflammatory microenvironments. Activated immune cells following brain damage are known to impact the proliferation, survival, migration, and differentiation of NSCs (Addington et al., 2015; Covacu & Brundin, 2017). Several studies have shown that an increase of M2 immune cells after brain injury has neurogenic effects on NSCs *in vivo* and *in vitro* (Choi et al., 2017; Vay et al., 2018). Accordingly, we suggest the possibility that ES enhances the anti-inflammatory state of the lesion environment, leading to increased survival and proliferation of NSCs.

However, while ES treatment showed a significant suppression of cortical neuronal die-off (evidenced by sustained NeuN expression at the perilesional cortex as compared to untreated TBI group), this effect was also observed in the sham group (Figure 7b). These results suggest that the electrode implantation into the cortical lesion alone, regardless of ES, could improve the recovery of the brain function by increasing neuronal viability in the cortex after seven days post-TBI. We also observed more M2 microglia in the groups with an electrode implantation, which may imply a possible involvement of M2 microglia in the neuronal viability after TBI.

In conclusion, to the best of our knowledge, our study is the first to demonstrate the overall impact of ES during the acute phase of TBI recovery on the promoted pro-healing phenotypes of microglia, the increased population of NSCs and Nestin^+^ astrocytes, and the enhanced viability of cortical neurons. Based on our findings, we propose a plausible scenario for intercellular communication among microglia, NSCs, astrocytes, and neurons (Figure 9). As illustrated in Figure 9a, after TBI, pro-healing immune cells and NSCs were confined near the edge of the cortical lesion, not being able to extend their protective influence over the entire perilesional region during the recovery process. When ES was applied, the effective region of occupancy by pro-healing immune cells (especially microglia) and Nestin^+^ cells (derived from either NSCs or astrocytes) penetrated deeper into the distant perilesional cortex. We confirmed enhanced neuronal viability after TBI over the broader perilesional cortex, which is believed to be accompanied by changes in injury-associated cellular and inflammatory milieu. Figure 9b illustrates the emergence of various types of Nestin^+^ cells by ES that include Nestin^+^ pericyte, Nestin^+^ NSC, and Nestin^+^GFAP^+^ NSC. However, neither the origin of the emergent NSCs nor the mechanism of functional alterations by ES is known, and it will be an intriguing research topic to further investigate these mysteries. Additionally, for effective therapeutic application of ES, optimization of ES parameters, including frequency, magnitude, duration of stimulation, and number of treatments, for overall neural tissue regeneration should be thoroughly explored. Nevertheless, our results strongly support the potential benefit of the therapeutic use of ES during the acute phase of TBI to regulate neuroinflammation and to enhance neuroregeneration for improved healing.

**FIGURE 9.**
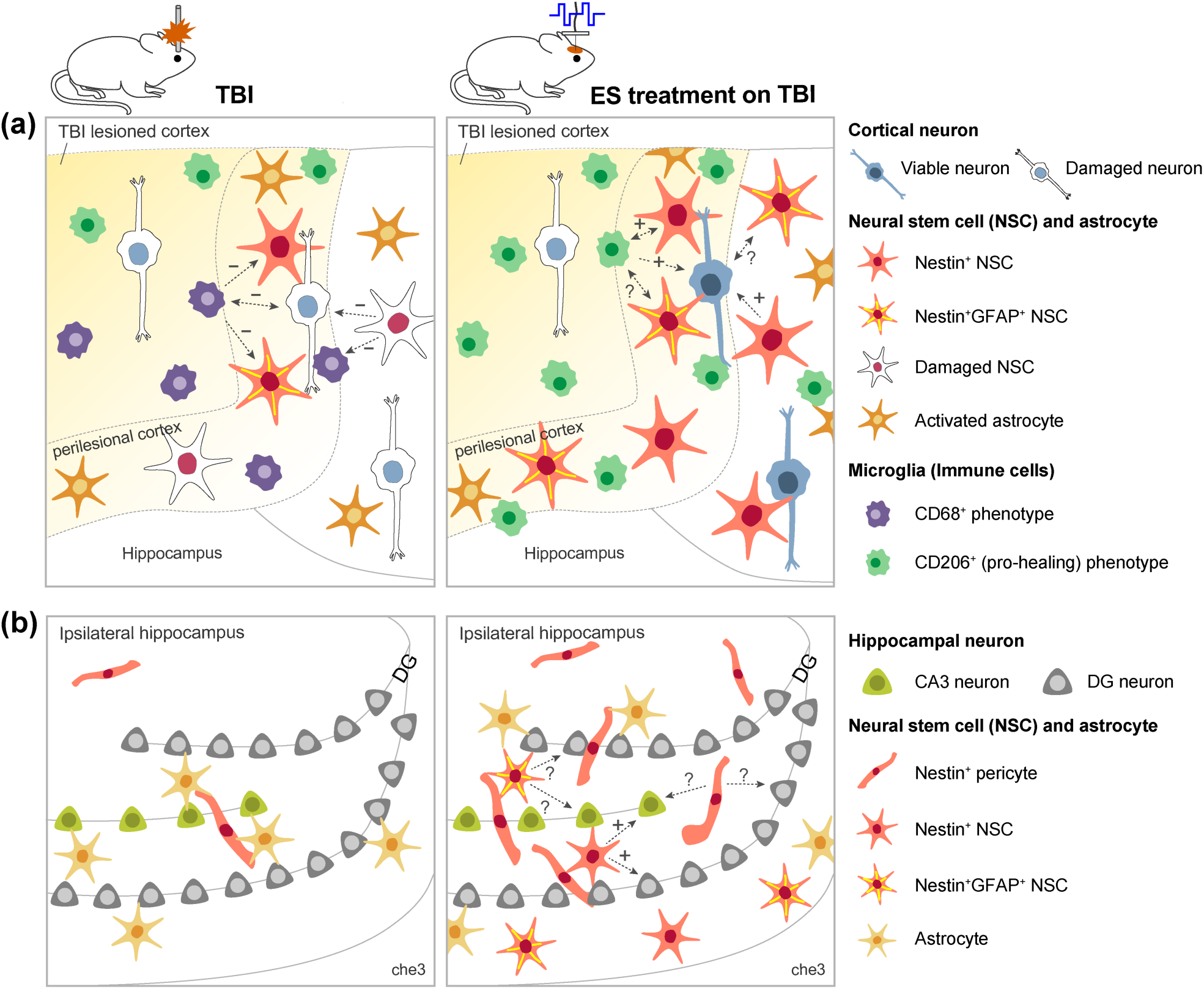
Graphical abstract showing a plausible scenario for intercellular communication among microglia, neural stem cells, and neurons in untreated TBI and ES groups. The expected influence of cell 1 on cell 2 is indicated in the direction of the dotted arrow. Next to the arrow, supportive or detrimental influences predicted based on literature were indicated by an “+” or “-”, and unknown interactions were indicated by a question mark. (a) and (b) show perilesional cortex and ipsilateral hippocampus, respectively.

## Author Contributions

All the authors read and approved the final manuscript. *Conceptualization*, E.P., J.G.L., J.H.S., and R.V.B.; *Data Curation*, E.P.; *Formal Analysis*, E.P., and J.G.L.; *Investigation*, E.P., M.A.-V., and M.I.B.; *Methodology*, E.P., J.G.L., M.A.-V., M.I.B., and N.M.; *Project Administration*, E.P., J.G.L., J.H.S., and R.V.B.; *Validation*, E.P., M.A.-V., and M.I.B.; *Writing – Original Draft*, E.P.; *Writing – Review & Editing*, E.P., J.G.L., N.M., J.H.S., and R.V.B.; *Funding Acquisition*, J.H.S, and R.V.B.; *Resources*, J.H.S., and R.V.B.; *Software*, E.P., and J.G.L., *Supervision*, J.G.L., J.H.S., and R.V.B.

## Acknowledgments

This paper is based on a research which has been conducted as part of the KAIST-funded Global Singularity Research Program for 2020. This work was supported by the National Institute of Neurological Disorders and Stroke (NINDS), of the National Institutes of Health (NIH), R01NS079739-05.

## Data Accessibility Statement

All data needed to evaluate the conclusions in the paper are present in the paper and/or the Supporting Information. Additional data related to this paper may be requested from the authors.

## Supporting Information for

### Figures

**FIGURE S1.**
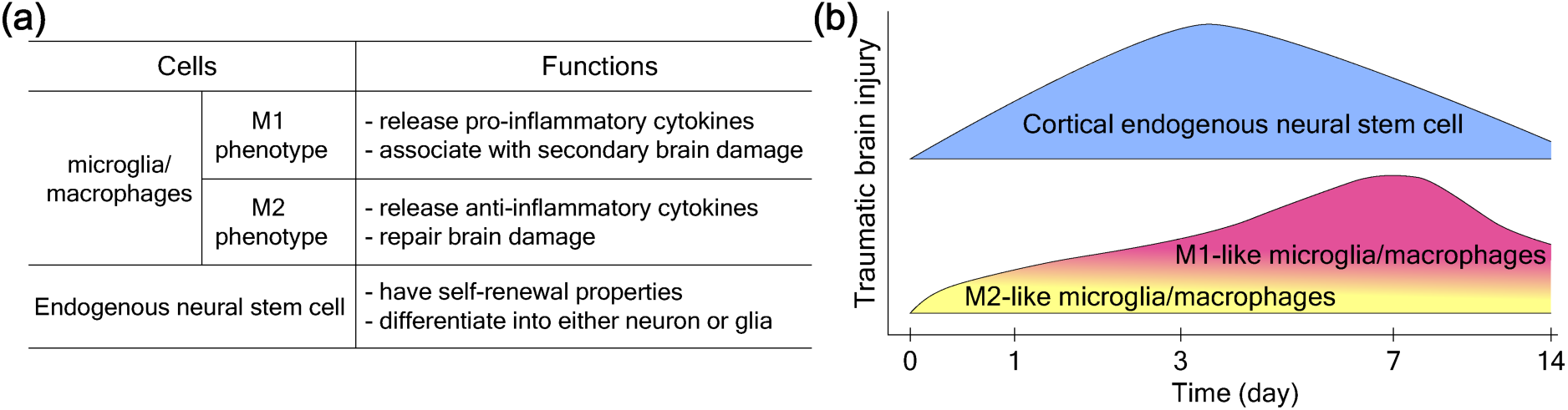
Cellular response in early phase of TBI. (a) Function of microglia/macrophage subtypes and endogenous neural stem cells (NSCs) following TBI. (**B**) Temporal changes in inflammatory responses and NSCs after TBI. Within 3 days after TBI, mixed M1/M2 phenotype of microglia/macrophages appears, then M1-phenotype predominates (Kumar, Alvarez-Croda, Stoica, Faden, & Loane, 2016; Simon et al., 2017; Wang et al., 2013). Endogenous NSCs appear following injury onset, peaking at 3 days post-TBI then decreasing (Itoh, Satou, Hashimoto, & Ito, 2005; Yi et al., 2013). Temporally, these immune and neural stem cell responses overlap.

**FIGURE S2.**
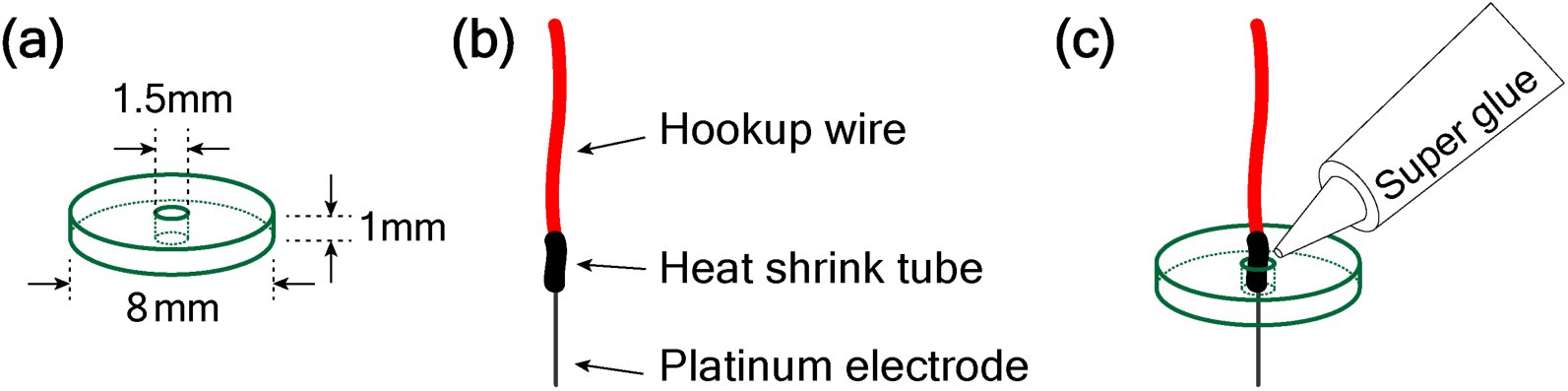
Illustration of an implantable ES device used in this study. (c) PDMS block: Prepolymer Sylgard 184 and its curing agent were cast at a height of 1mm in a ratio of 10:1. Next, the block was cut with a punch of 8mm in diameter. A 1.5mm punch was used in the center of the block to create a hole to insert the electrode. (d) Platinum electrode and hookup wire were connected by soldering and then encase them with a heat shrink tube. (e) Electrodes were placed in the hole of the PDMS block then affixed with super glue.

**FIGURE S3.**
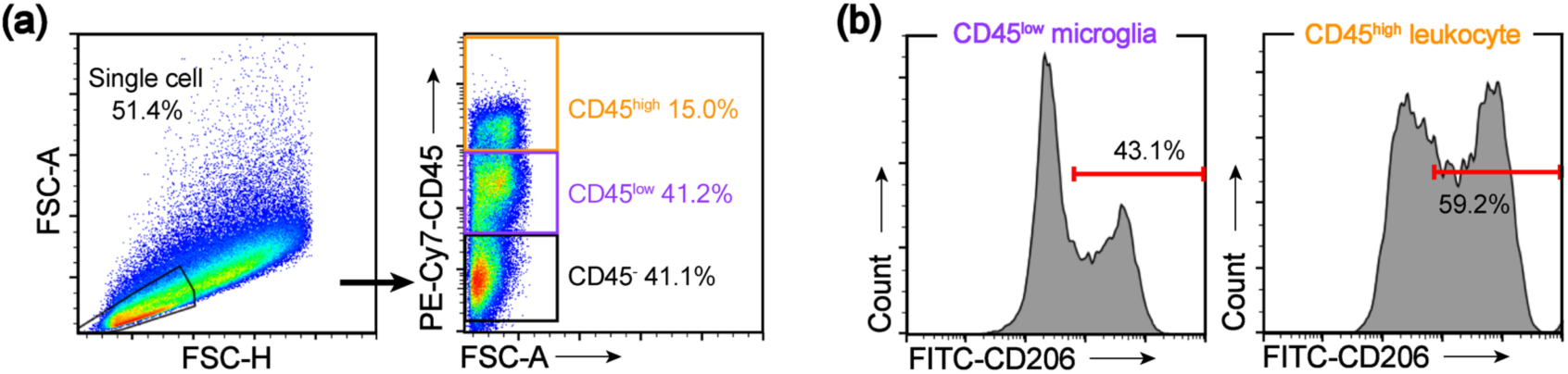
Representative gating strategy used to identify CD206^+^ immune cell subsets in perilesional brain tissue by flow cytometry. (a) Based on CD45 intensity, single cells were distinguished into CD45^low^ microglia and CD45^high^ leukocytes population. (b) CD206^+^ population was further gated to identify the subsets.

**FIGURE S4.**
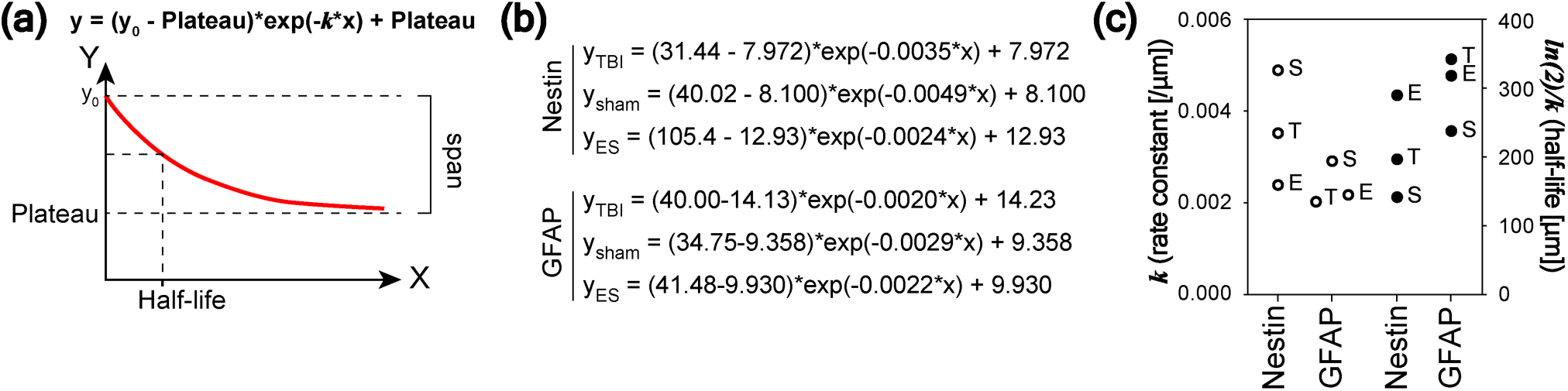
One-phase exponential decay curve and fitting results. (a) Illustration of one-phase exponential decay curve and its equation, shown on the graph. In this study, the x-axis was for a distance from the perilesional rims, and the y-axis was for a mean intensity of Nestin or GFAP as shown in Figure 5c and 5d. y_0_ is the initial intensity value when x is zero. The value *k* in the equation represents the rate of intensity decrease over a distance. Half-life (*ln(2)/k*) is a distance required for intensity to reduce to 50 % of span. (b) Fitted function for the mean intensity of Nestin and GFAP in TBI, sham, and ES groups. (c) Fitted values of rate constant and half-life.

